# Spectral slope and Lempel-Ziv complexity as robust markers of brain states during sleep and wakefulness

**DOI:** 10.1101/2022.09.10.507390

**Authors:** Christopher Höhn, Michael A. Hahn, Janna D. Lendner, Kerstin Hoedlmoser

## Abstract

Spectral slope and Lempel-Ziv complexity are affected in many neurophysiological disorders and are modulated by sleep, anesthesia, and aging. Yet, few studies have explored the relationship between these two parameters. We evaluated the impact of sleep stage and task-engagement (resting, attention and memory) on spectral slope and Lempel-Ziv complexity in a narrow- (30 – 45Hz) and broadband (1 – 45Hz) frequency range in 28 healthy males (21.54 ± 1.90 years) over three recordings. Only in the broadband range, the slope steepens and complexity decreases continuously from wakefulness to N3. However, REM sleep is best discriminated by the narrowband slope. Importantly, slope and complexity also differentiate between tasks during wakefulness. While the narrowband complexity decreases across tasks, the slope is flattening with task engagement in both frequency ranges. In general, broadband slope and complexity are strongly positively correlated, but we observe a dissociation between them in the narrowband range. Critically, only the narrowband slope is associated with better Go/Nogo task performance. Our results demonstrate that slope and complexity are both powerful indices of sleep depth, task engagement and cognitive performance. While the broadband range is better suited to discriminate between brain states, especially the narrowband slope is a unique marker of task performance.

## Introduction

To date, neural oscillations are still the most prominent electrophysiological signature of human brain activity. For instance, wakeful resting is typically characterized by pronounced alpha-band activity (8 – 12Hz), which is suppressed in active task engagement (Kirstein, 2007; Klimesch et al., 1993; Klimesch, 1999). During sleep, different stages are best described by characteristic oscillatory events like sleep spindles and slow oscillations (Davis et al., 1938; Richard et al., 2012; Terzano et al., 2002). However, recent evidence suggests that irregular and aperiodic brain activity, measured by Lempel-Ziv complexity (Lempel & Ziv, 1976 and Welch, 1984) and the spectral exponent *ß* (i.e., the magnitude of decay in power with increasing frequency; He, 2014), also carries meaningful information about different brain states that might track varying arousal levels. Specifically, the spectral exponent has been discussed as a marker of the brain’s excitation and inhibition (E/I) balance (Gao et al., 2017), which is impaired in a variety of clinical conditions such as the attention deficit hyperactivity disorder (ADHD, Karalunas et al., 2022; Robertson et al., 2019), autism (Gao & Penzes, 2015; Rubenstein & Merzenich, 2003) and epilepsy (Symonds, 1959; Wong, 2010). In addition, epilepsy has further been associated with alterations in neural complexity (Aarabi & He, 2012; Zhu et al., 2017).

Conceptually, spectral exponent and Lempel-Ziv complexity (frequently used as an estimate of neural complexity) are derived by different analytical approaches from the underlying electrophysiological signal. The spectral exponent is a measure obtained from the frequency domain, while Lempel-Ziv complexity is computed in the time domain. Lempel-Ziv complexity (Welch, 1984) reflects the regularity and compressibility of a signal in time-domain (Lau et al., 2022) and is strongly influenced by oscillatory activity (González et al., 2022; Tosun et al., 2019), with highly regular signals resulting in low complexity whereas unpredictable, irregular signals lead to high complexity (see Figure 1 – Figure Supplement 1). In contrast, the spectral exponent reflects the absolute value of the slope (i.e., steepness) of a signal’s power spectrum in frequency-domain, which is thought to be mainly aperiodic and therefore irregular (Donoghue et al., 2020). Thus, more aperiodicity in the signal is accompanied by both, a flatter (i.e., less negative) slope and higher complexity. However, additional oscillatory components added to the signal should only decrease the complexity value with the slope remaining relatively unaffected. In the following, we always use and refer to the spectral slope instead of the spectral exponent (which would be the absolute value of the slope) to avoid any ambiguity due to different terms that are common to describe the same parameter (e.g., 1/f signal and scale free or aperiodic activity).

**Figure 1.**
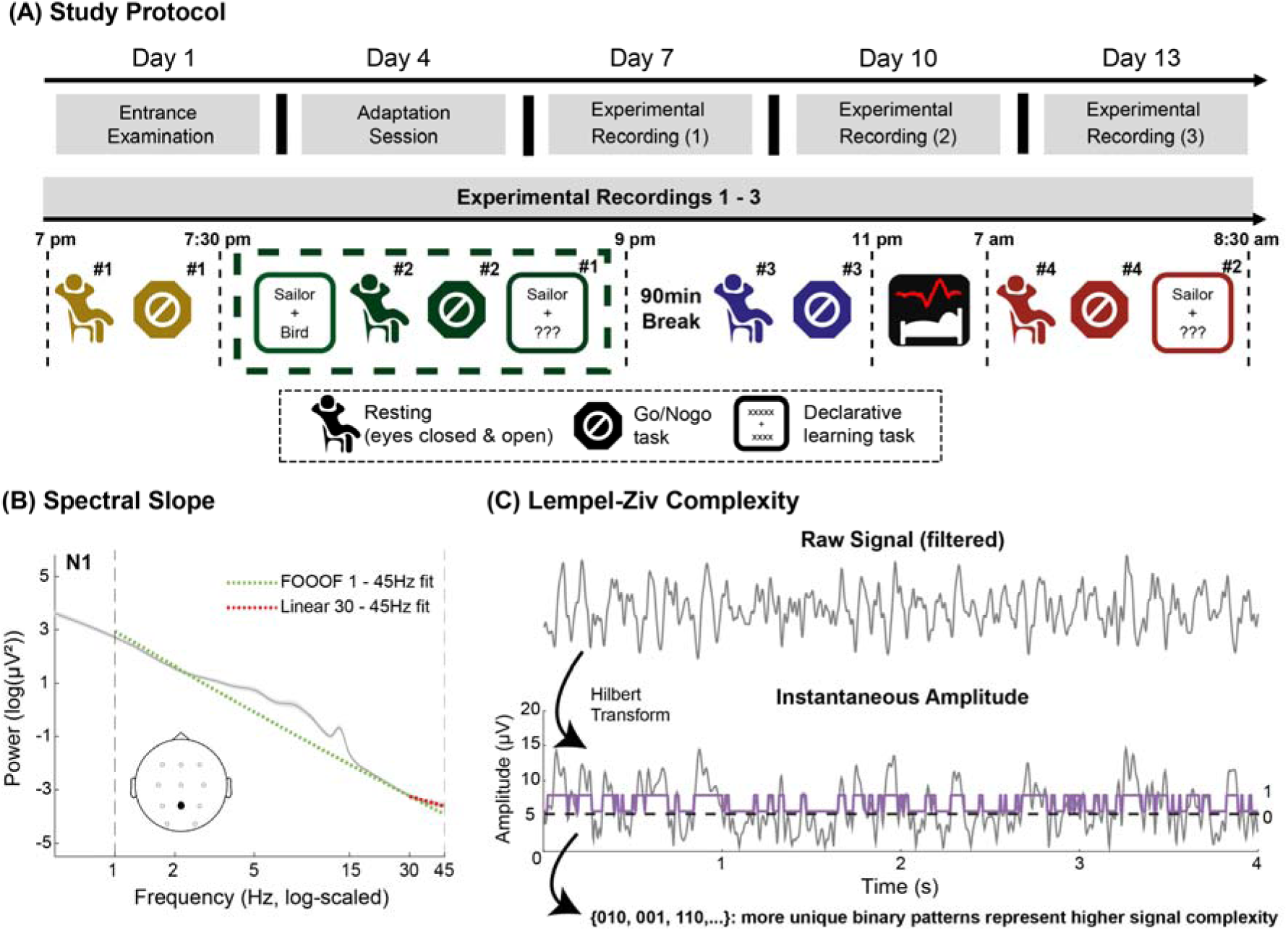
(A): Overview of the experimental protocol. EEG was recorded throughout all tasks and during sleep (with full-night polysomnography) on the experimental days 7, 10 and 13. The tasks, which are highlighted by a dashed, dark-green rectangle were primarily used to analyze the effects of engagement in different cognitive tasks during wakefulness. The adaptation night only served familiarization purposes and was not included in any of the analyses. Results from the entrance examination questionnaires are presented in Supplementary file – Table 1. (B): Example of the spectral slope estimation during N1 sleep. For illustration purposes, data is shown for the electrode Pz averaged over all subjects and sleep recordings. The spectral slope was fitted within 1 – 45Hz (broadband, dashed green line) and 30 – 45Hz (narrowband, dashed red line). (C): Schematic overview of the Lempel-Ziv complexity calculation based on a random 4s epoch from electrode Pz of a subject during resting with closed eyes. First, the raw signal, filtered within a certain frequency range, is Hilbert transformed. Second, the resulting data is binarized around its median amplitude and stored as a vector of zeros and ones. Lastly, the Lempel-Ziv-Welch algorithm (Welch, 1984) is applied on this binary sequence in order to obtain a complexity value, which is driven by the number of unique repetitions of ones and zeros. **Figure 1: Supplement 1.** Illustration of the effect of signal regularity on resulting Lempel-Ziv complexity values and the shape of their power-spectra. The complexity values increase from a binary boxcar signal (purple) to a pure 10 Hz alpha oscillation (blue) and further to the same oscillation with additional pink noise (orange). Pure pink noise (yellow) has the highest complexity.

Although the spectral slope is derived from the frequency domain and Lempel-Ziv complexity from the time domain, the literature suggests that both metrics capture changes in brain state in a surprisingly similar way, despite their apparent analytical differences. Regarding arousal levels, multiple studies showed that the spectral slope steepens (i.e., becomes more negative) from wakefulness to anesthesia (Colombo et al., 2019; Gao et al., 2017; Lendner et al., 2020; Waschke et al., 2021), while others showed the same pattern for Lempel-Ziv complexity, which also decreases from wakefulness to anesthesia (Ferenets et al., 2007; Zhang et al., 2001). Similarly, spectral slope (Lendner et al., 2020; Ma et al., 2018; Miskovic et al., 2019; Pereda et al., 1998) and Lempel-Ziv complexity (Andrillon et al., 2016; Schartner et al., 2017) both decrease with increasing sleep depth (i.e., from wakefulness to N3 sleep). Besides changes in arousal, recent evidence from Waschke et al. (2021) further suggests that the spectral slope also tracks the level of attention, whereby higher levels of attention and quicker response times are indexed by flatter slopes during wakefulness. This is in line with findings from other studies, which showed that the slope is indicative of cognitive processing speed (Ouyang et al., 2020; Pathania et al., 2022) and modulated by cognitive decline in ageing (Dave et al., 2018; Voytek et al., 2015; Voytek & Knight, 2015). Similarly, higher Lempel-Ziv complexity values also relate to faster reaction times on a trial-by-trial basis, thus likewise serving as a proxy of attention or processing speed (Mediano et al., 2021). Following these commonalities in the modulation of the spectral slope and Lempel-Ziv complexity by brain state changes through sleep, anesthesia and task-demand, it might be inferred that these two parameters may be indicative of similar neural processes. Yet, there is only limited research examining the potential relationship between these parameters (Medel et al., 2020).

With respect to the influence of different frequency contents on the estimation of the spectral slope and Lempel-Ziv complexity, no optimal frequency settings are established yet for any of the two parameters. The heterogeneity of frequency content on which the calculations of both measures are based might be responsible for some disparate results in the current literature, thus further hampering our understanding of the contribution of aperiodic brain activity to healthy brain functioning. For instance, González et al. (2022) suggest that particularly for complexity, lower frequencies (≤ 12Hz) are more informative than higher frequencies when differentiating between sleep and wakefulness. For the estimation of the spectral slope, researchers have argued either in favor of broadband (Karalunas et al., 2022; Podvalny et al., 2015; Waschke et al., 2021) or narrowband (Gao et al., 2017; Lendner et al., 2020) frequency ranges, commonly within a range between 1 – 45/50Hz to avoid strong bends (i.e., knees) in the power spectrum below or above these limits. While broadband ranges (e.g., 1 – 45Hz) encompass more of the total signal power and result in better overall slope-fits (Donoghue et al., 2020; Gerster et al., 2022), narrowband ranges (e.g., 25/30 – 45Hz) are less affected by low-frequency oscillatory activity and are therefore reflecting mostly pure aperiodic activity (Gao et al., 2017; Lendner et al., 2020).

Taken together, the slope and complexity research findings suggest a functional overlap of Lempel-Ziv complexity and spectral slope in tracking different brain states. However, to date a direct comparison of these two measures across different brain states during sleep and wakefulness is missing. Thus, the relationship between slope and complexity across brain states still remains unclear as it has only been compared between rested wakefulness and anesthesia so far (Medel et al., 2020). Additionally, little is known about the sensitivity of the spectral slope and Lempel-Ziv complexity to changes in brain activity during wakefulness in general. As potential markers of arousal and attention, the two parameters might likely be also affected by task engagement and different cognitive tasks in general, which require varying cognitive resources. Finally, it is unclear how the two measures are affected by selecting different frequency contents for their calculation.

Here, we leverage an expansive, within-subject design with multiple sleep and wake recordings to investigate (1) whether spectral slope and Lempel-Ziv complexity are modulated by different brain states during sleep and wakefulness and (2) to what extent they are related to each other as well as their functional significance for cognition. Using multiple recordings per subject, we try to overcome a limitation of most previous research that only relies on single session recordings, thus, limiting insights into the robustness of the observed effects. First, we assess the performance of the spectral slope and Lempel-Ziv complexity in delineating sleep from wakefulness. Second, we investigate the influence of task-engagement in different cognitive tasks on both measures by using simple resting sessions, an auditory attention (Go/Nogo) and a declarative memory task. Third, we analyze the relationship between the spectral slope and Lempel-Ziv complexity across sleep stages and tasks using either narrow-or broadband frequency ranges for estimation. Finally, we probe whether the two parameters can track behavioral performance in the Go/Nogo and declarative memory tasks.

## Results

We utilized data from a recently published study (Höhn et al., 2021; Schmid et al., 2021) that investigated the effects of different light conditions on alertness, sleep, and memory consolidation. The subjects underwent the same experimental protocol on three different days under highly controlled and standardized light conditions. On three consecutive experimental nights, multiple tasks were conducted before and after sleep, including two resting sessions with either eyes closed or open, an auditory attention task (Go/Nogo) and a declarative memory task (cf., Figure 1A). We calculated spectral slopes and Lempel-Ziv complexity for all sleep stages and tasks in a narrow-(30 – 45Hz) and broadband (1 – 45Hz) frequency range (cf., Figure 1B and C). We decided to set the upper frequency limit to 45Hz to avoid any influences of line-noise and the need of fitting a knee in higher frequencies. Following the results from Lendner et al. (2020) and Lendner et al. (2022), we restricted the narrowband range to 30 – 45Hz as this was shown to closely track the hypnogram and has also been used as a proxy for excitation/inhibition balance in the brain (Gao et al., 2017).

### Spectral slope and Lempel-Ziv complexity delineate brain states during sleep

First, we strived for replicating previous findings, which showed that sleep stages could be differentiated solely based on the spectral slope and signal complexity. The effect of sleep stage was assessed for the spectral slope and Lempel-Ziv complexity in each frequency range (30 – 45Hz and 1 – 45Hz) with semi-parametric Wald-Type Statistics (WTS; Friedrich et al., 2019) averaged over all electrodes while considering the three repeated measurements.

The narrowband (30 – 45Hz) slope (*WTS* (4) = 133.57, *p* <.001) and complexity (*WTS* (4) = 21.64, *p* =.004) models both indicated significant modulations by sleep stage. In line with previous research, the narrowband slope was significantly steeper in all sleep stages compared to wakefulness with the steepest slope during REM sleep. In contrast, the narrowband complexity slightly increased from wake to sleep and showed a diverging pattern in comparison to the narrowband slope (see Figure 2). To test whether the results in this frequency range (30 – 45Hz) might be driven or explained by changes in muscular activity (i.e., EMG), we also computed the slope and complexity of the EMG channels (see Figure 2 – Figure Supplements 1 & 2). Especially during sleep, the EMG did not account for a significant amount of variation of the EEG slope or complexity across sleep stages. In wakefulness, the EMG slope and complexity also varied between tasks and were especially affected by the learning task, which involved verbal communication and therefore naturally elevated EMG activity. However, when partialling out the EMG from the EEG data, the modulation of the EEG slope and complexity by tasks remained largely unaffected, indicating that EMG activity did not confound our results.

**Figure 2.**
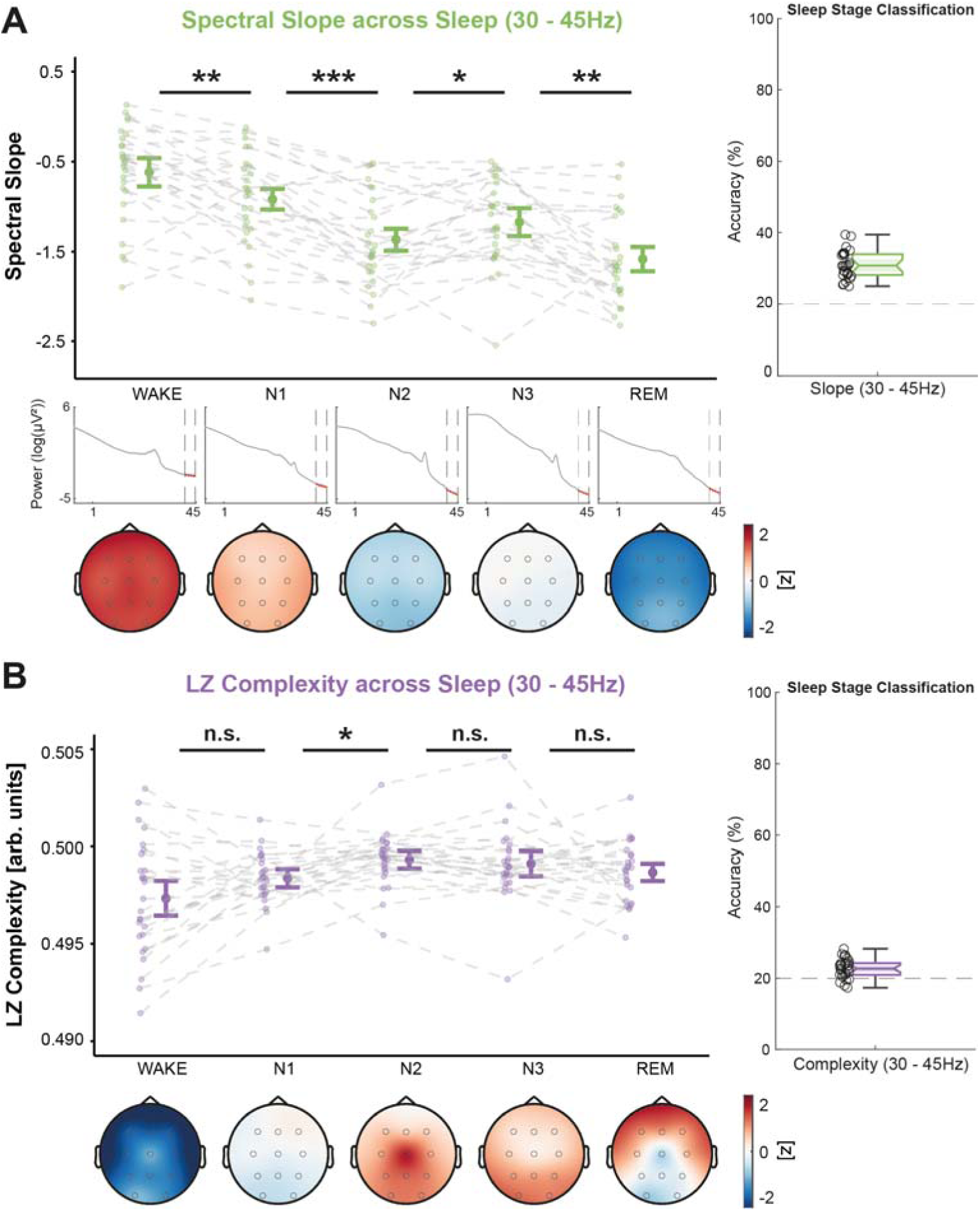
Spectral slope (green, A) and Lempel-Ziv (LZ) complexity (purple, B) from 30 – 45Hz across sleep, averaged over all lab-sessions per subject. Center figures show the data averaged over all electrodes and topographical maps are provided below (color-coding refers to z-values of slope or complexity computed from the grand average across all sleep stages). In A, the log-log power spectra are provided for each sleep stage to illustrate the slope changes across different sleep stages. Classification accuracies are shown on the right-hand side. A: The spectral slope decreases from wakefulness across all sleep stages to REM sleep with a small temporary increase during N3 sleep. B: Lempel-Ziv complexity increases from shallow N1 to light N2 sleep and is in general less modulated by sleep stage than the spectral slope. ***: *p* <.001, **: *p* ≤.010, *: *p* ≤.050, n.s.: *p* >.050; all p-values are adjusted for multiple comparisons; error-bars represent 95% confidence intervals (*N* = 27). **Figure 2: Supplement 1.** Control analyses including the narrowband spectral slope from the EMG. A: The negative correlations between EEG slope and sleep stage do not change when partialling out the EMG slope. B: While the average EEG slope is negatively correlated with sleep stage, the EMG slope is even slightly positively correlated with sleep stage and significantly different from the EEG slope correlation. C: The positive correlations between EEG slope and the cognitive tasks (ordered ascendingly regarding their slope) are not diminished when controlling for the EMG. D: While the correlation between the EMG slope and the tasks is slightly higher than between the EEG slope and the tasks, partialling out the EMG from the EEG slope does not significantly reduce the correlation. E & F: Differential modulation of the EEG & EMG slopes across sleep stages and tasks. **Figure 2: Supplement 2.** Control analyses including the narrowband Lempel-Ziv complexity (LZC) from the EMG. A: The positive correlations between EEG complexity and sleep stage do not change when partialling out the EMG complexity. B: While both, the average EEG and EMG complexity are positively correlated with sleep stage, the partial correlation controlling for EMG complexity does not shrink substantially. C: The negative correlations between EEG complexity and the cognitive tasks are not changed substantially by partialling out the EMG. D: Both, the average EEG and EMG complexity are negatively correlated with the cognitive tasks but the partial correlation between EEG complexity and the tasks controlled for the EMG is not significantly smaller. E & F: Differential modulation of the EEG & EMG complexity across sleep stages and tasks.

When the broadband (1 – 45Hz) frequency range was used for estimation, the effect of sleep stage was much more pronounced in both parameters (spectral slope: *WTS* (4) = 1088.28, *p* <.001; Lempel-Ziv complexity: *WTS* (4) = 857.60, *p* <.001). Both, the broadband slope and complexity significantly decreased from shallow (N1) to deep NREM sleep (N3). For REM sleep, however, both markers increased again in remarkably similar ways (see Figure 3), arguably reflecting more wake-like brain activity in the broadband range.

**Figure 3.**
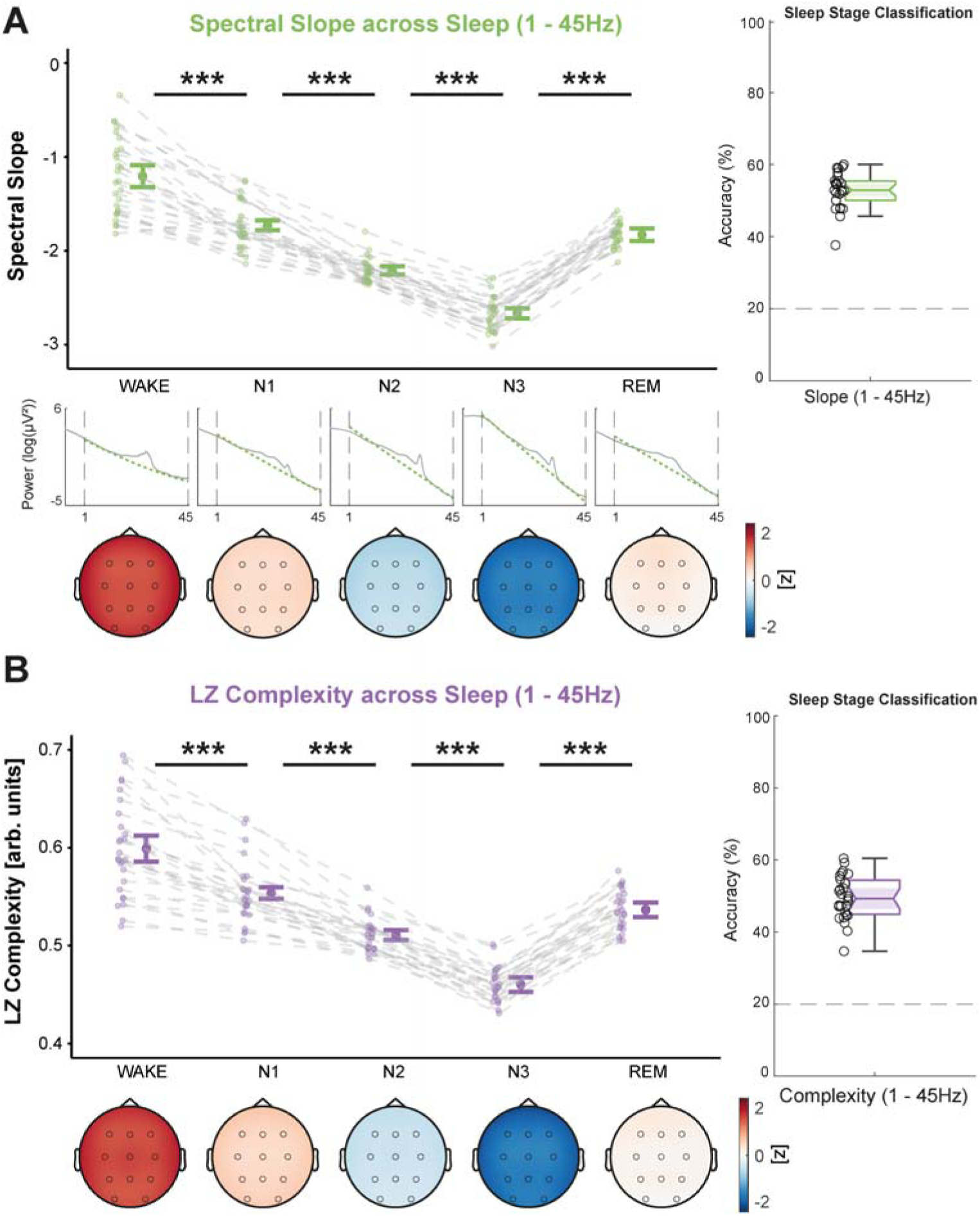
Spectral slope (green, A) and Lempel-Ziv (LZ) complexity (purple, B) from 1 – 45Hz across sleep, averaged over all lab-sessions per subject. Center figures show the data averaged over all electrodes and topographical maps are provided below (color-coding refers to z-values of slope or complexity computed from the grand average across all sleep stages). In A, the log-log power spectra for each sleep stage are provided to illustrate the broadband slope differences across sleep stages. Classification accuracies are shown on the right-hand side. A: Spectral slope steepens from wakefulness to N3 sleep but flattens to some extent in REM sleep. B: Lempel-Ziv complexity shows the same pattern as the spectral slope and likewise decreases from wakefulness to N3 with a subsequent increase in REM sleep. ***: *p* <.001, **: *p* ≤.010, *: *p* ≤.050, n.s.: *p* >.050; p-values are adjusted for multiple comparisons; error-bars represent 95% confidence intervals (*N* = 27).

We found no significant effects of the repeated measurements (all padj. ≥.419 after correcting for multiple comparisons), revealing that the effect of sleep stage robustly emerged in all individual recordings per subject. To evaluate the topographical distribution of the spectral slope and Lempel-Ziv complexity, we additionally ran a multivariate pattern analysis (MVPA) with multi-class linear discriminant analyses (LDA). With this MVPA, we quantified how well sleep stages could be decoded by taking the topographical distribution of the slope and complexity values into account. In both frequency ranges and for both parameters, classification accuracies were always significantly above chance level (20%, *p* <.001) and in general higher for the broadband (1 – 45Hz) than for the narrowband (30 – 45Hz) frequency range (*WTS* (1) = 724.34, *p* <.001). For both frequency ranges, the spectral slope was more informative about the underlying brain state (i.e., yielded higher classification accuracies) than Lempel-Ziv complexity (*WTS* (1) = 182.09, *p* <.001), especially in the narrowband range (spectral slope: 30.96%, Lempel-Ziv complexity: 22.67%; see Figure 4A).

**Figure 4.**
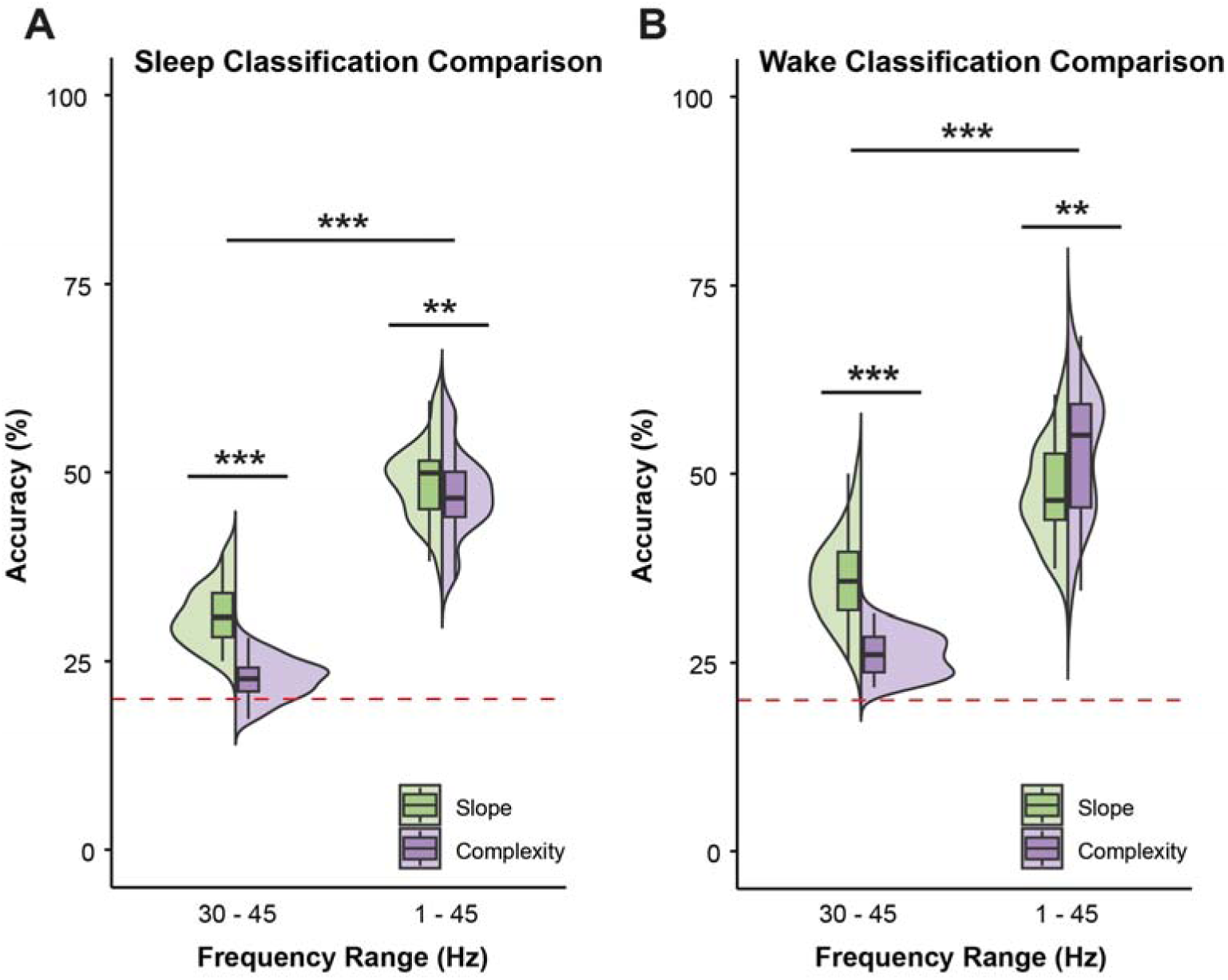
A: Direct comparison of the multi-class classification accuracies across sleep for the spectral slope and Lempel-Ziv complexity within 30 – 45Hz vs. 1 – 45Hz. Sleep stages could be decoded more precisely for both parameters in the broadband frequency range, but classification accuracy was significantly higher for the spectral slope, especially in the narrowband range. B: Comparison of the classification accuracies across tasks for slope and complexity in both frequency ranges. Again, the 1 – 45Hz range yielded better decoding performance in general, but the performance in the broadband range was significantly higher for Lempel-Ziv complexity than for the spectral slope, arguably due to the difference in complexity between the two resting conditions, which is not present in the slope. The red dotted line represents chance level (20% for 5 sleep stages or tasks). ***: *p* <.001, **: *p* ≤.010, *: *p* ≤.050, n.s.: *p* >.050.

During wakefulness, the MVPA results also indicated an above chance classification performance for all tasks (20%, *p* <.001). Similar to the sleep-results, classification accuracy was higher when using the broadband instead of the narrowband frequency range (*WTS* (1) = 364.18, *p* <.001). The spectral slope was again more informative in the narrowband range (Slope: 35.96%, Complexity: 25.98%, *WTS* (1) = 89.53, *p* <.001) while Lempel-Ziv complexity yielded better results in the broadband range (Slope: 47.66%, Complexity: 52.69%, *WTS* (1) = 15.12, *p* =.001; see Figure 4B).

### Spectral slope and Lempel-Ziv complexity vary across different tasks

Next, we investigated whether spectral slope and Lempel-Ziv complexity can differentiate between resting and task-engagement and might also track different cognitive tasks. We calculated both markers from resting sessions with eyes closed (REC) and eyes open (REO), an auditory Go/Nogo task (GNG), an encoding session (ENC) from a declarative memory task as well as its according retrieval session (RET). For these analyses, we focused on the task data from the evening recordings (see dashed dark-green rectangle in Figure 1A). Theoretically, a task-engagement effect, representing a shift towards excitation instead of inhibition (i.e., flatter slopes and higher complexity), should be visible between the resting sessions and the Go/Nogo or learning task. Since the Go/Nogo task was mainly auditory and should rely on different cognitive resources than the more visual/verbal declarative learning task, we also expected differences between the Go/Nogo, encoding and retrieval sessions.

In the narrowband frequency range (30 – 45Hz), we observed a significant flattening (i.e., values closer to zero) of the slope (*WTS* (4) = 56.64, *p* <.001) but a decrease in complexity (*WTS* (4) = 199.55, *p* <.001) from resting sessions to active tasks (see Figure 5). As expected, the spectral slope was flatter during the Go/Nogo (GNG vs. REC: *WTS* (1) = 21.05, *p_adj._* <.001; GNG vs. REO: *WTS* (1) = 16.53, *p_adj._* =.001), encoding (ENC vs. REC: *WTS* (1) = 20.73, *p_adj._* <.001; ENC vs. REO: *WTS* (1) = 15.56, *p_adj._* =.001) and retrieval tasks (RET vs. REC: *WTS* (1) = 48.84, *p_adj._* <.001; RET vs. REO: *WTS* (1) = 41.37, *p_adj._* <.001) as compared to resting. However, there was also an additional flattening of the narrowband slope present during the retrieval task in comparison to the two other tasks (RET vs. GNG: *WTS* (1) = 6.44, *p_adj._* =.021; RET vs. ENC: *WTS* (1) = 13.66, *p_adj._* =.001). Narrowband Lempel-Ziv complexity did not differ between the resting and Go/Nogo sessions (all *p_adj._* >.110) but decreased from the Go/Nogo to the encoding session (GNG vs. ENC: *WTS* (1) = 16.64, *p_adj._* <.001) and was lowest during retrieval (RET vs. GNG: *WTS* (1) = 98.74, *p_adj._* <.001, RET vs. ENC: *WTS* (1) = 31.11, *p_adj._* <.001).

**Figure 5.**
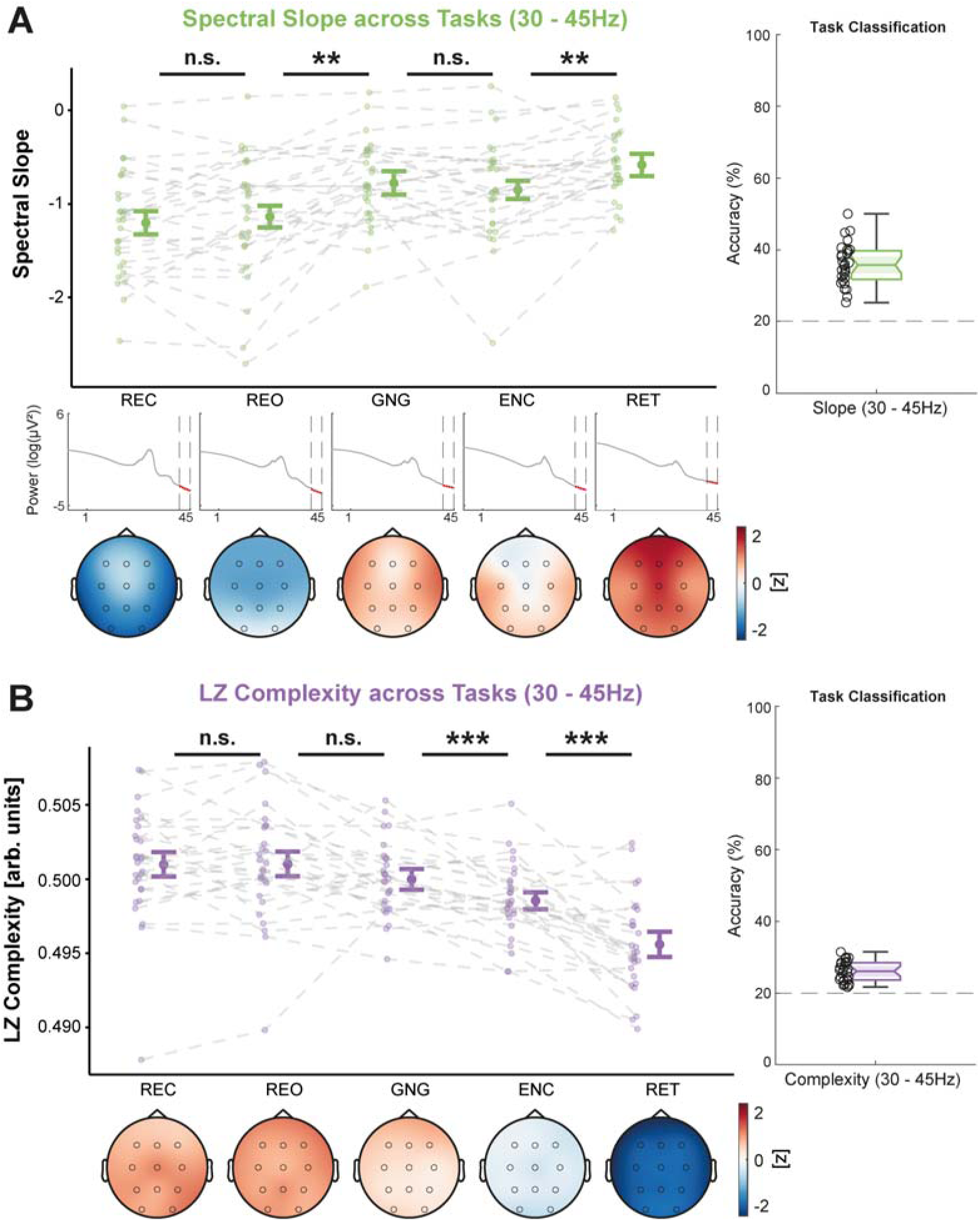
Spectral slope (green, A) and Lempel-Ziv (LZ) complexity (purple, B) from 30 – 45Hz across tasks, averaged over all lab-sessions per subject. Center figures show the data averaged over all channels and topographical maps are provided below (color-coding refers to z-values of slope or complexity computed from the grand average across all tasks). In A, the log-log power spectra for each task are provided to illustrate narrowband slope differences across tasks. Classification accuracies are shown on the right-hand side. A: The spectral slope flattens when engaging in cognitive tasks (Go/Nogo and learning) and is flattest during the retrieval session of the learning task. B: Lempel-Ziv complexity decreases with task-engagement and reaches its minimum during the retrieval session. ***: *p* <.001, **: *p* ≤.010, *: *p* ≤.050, n.s.: *p* >.050; p-values adjusted for multiple comparisons; error-bars show 95% confidence intervals (*N* = 28). **Figure 5: Supplement 1.** Slope and complexity (30 – 45Hz) across tasks averaged over all timepoints. **Figure 5: Supplement 2.** Slope and complexity (30 – 45Hz) across tasks using a different task-order (REC#1, GNG#1, ENC, REO#2, RET#1 instead of ENC, REC#2, REO#2, GNG#2, RET#1, cf., Figure 1).

When investigating the broadband frequency range (1 – 45Hz), we found that the diverging pattern between spectral slope and Lempel-Ziv complexity disappeared and both parameters were increasing (i.e., higher complexity values and flatter slopes indexed by less negative values) from resting sessions to active task engagement and were highest during the retrieval task (Slope: *WTS* (4) = 40.46, *p* <.001; Complexity: *WTS* (4) = 46.24, *p* <.001; see Figure 6). Within 1 – 45Hz, only Lempel-Ziv complexity additionally differed between the two resting sessions (eyes closed and eyes open), which is likely reflecting a difference in alpha power (8 – 12Hz). This offers further support to the notion of a greater influence of oscillatory signal components on estimates of complexity in contrast to the spectral slope as the major difference between resting with closed and open eyes in the power spectrum is a difference in alpha power. Again, we did not observe any effects of the repeated measurements (all *p_adj._* ≥.222).

**Figure 6.**
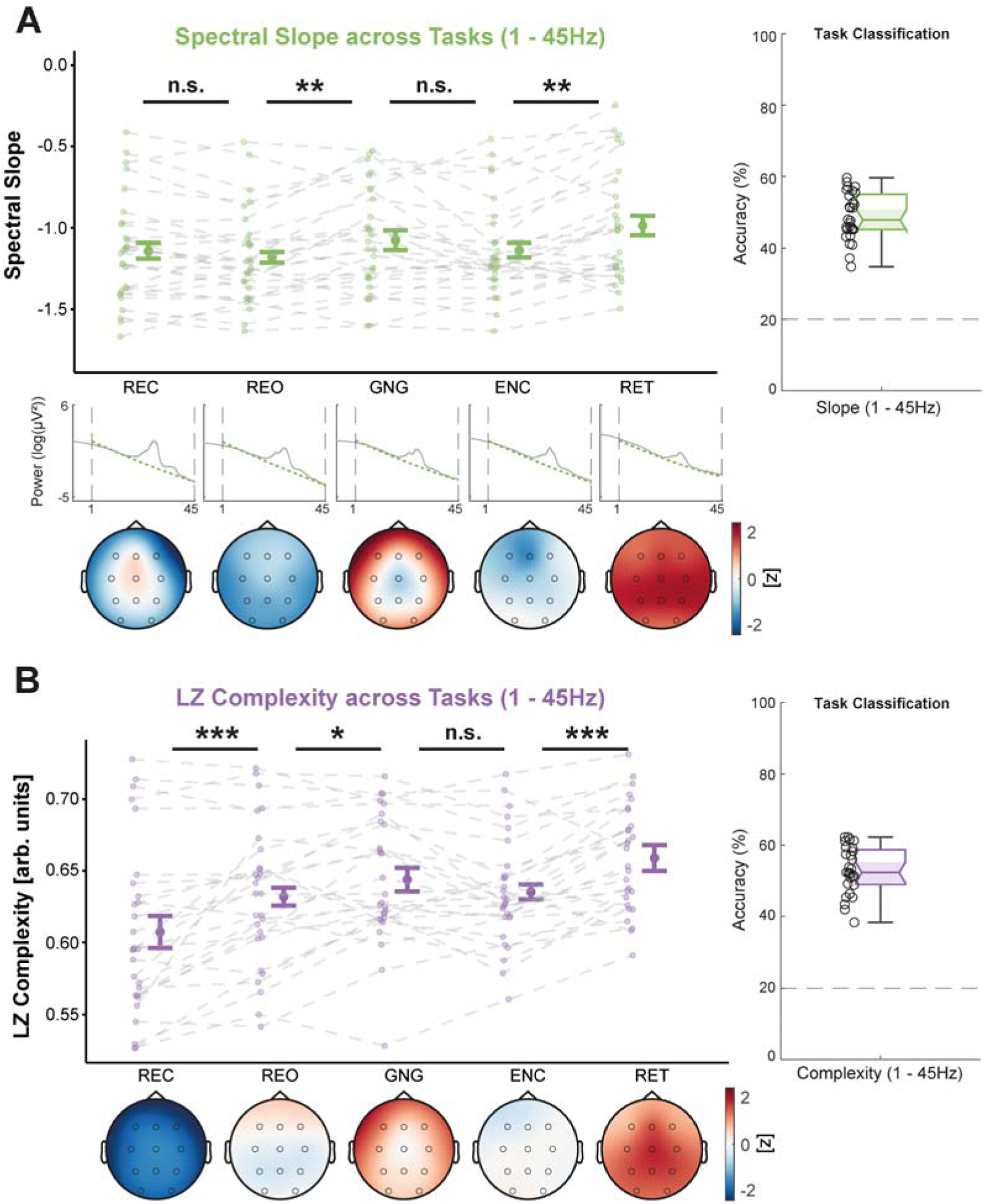
Spectral slope (green, A) and Lempel-Ziv (LZ) complexity (purple, B) from 1 – 45Hz across tasks, averaged over all lab-sessions per subject. Center figures show the data over all channels and topographical maps are provided below (color-coding refers to z-values of slope or complexity computed from the grand average over all tasks). In A, the log-log power spectra for each sleep stage are provided to illustrate broadband slope differences across tasks. Classification accuracies are shown on the right-hand side. A: The slope flattens from resting to the Go/Nogo task and is flattest during retrieval. B: Complexity increases from resting with closed to open eyes and is further elevated in all active tasks, peaking during retrieval. ***: *p* <.001, **: *p* ≤.010, *: *p* ≤.050, n.s.: *p* >.050; p-values adjusted for multiple comparisons; error-bars show 95% confidence intervals (*N* = 28). **Figure 6: Supplement 1.** Slope and complexity (1 – 45Hz) across tasks averaged over all timepoints. **Figure 6: Supplement 2.** Slope and complexity (1 – 45Hz) across tasks using a different task-order (REC#1, GNG#1, ENC, REO#2, RET#1 instead of ENC, REC#2, REO#2, GNG#2, RET#1, cf., Figure 1). **Figure 6: Supplement 3.** Slope and complexity between active tasks (GNG, ENC, RET) after correcting for the resting eyes open condition as baseline.

To control whether the results were confounded by the task order and thus solely reflect an increase in exhaustion or decrease in motivation, we repeated the analyses with the task data averaged over all available time points, which were not equal for all tasks (cf., Figure 1A, i.e., in this scenario the resting and Go/Nogo data was averaged over four different time points, the retrieval data over two time points and the encoding was only done once). We also repeated the analyses with a different order of tasks, resulting in “resting with eyes closed, Go/Nogo, encoding, resting eyes open and retrieval” instead of “encoding, resting with eyes closed, resting with eyes open, Go/Nogo and retrieval”. Both control analyses confirmed the same pattern as in the original analyses with a similar flattening of the broad-and narrowband slopes across tasks and an increase in broadband but a decrease in narrowband complexity (see Figure 5 – Figure Supplements 1 & 2 and Figure 6 – Figure Supplements 1 & 2).

An overview of the pairwise classification accuracies for all sleep stages and task pairings is presented in Supplementary file – Tables 2 and 3. All tasks and sleep stages could be differentiated above chance-level (50% in this context). As described above, the classification accuracy was in general higher for the broadband than the narrowband frequency range. However, in the narrowband frequency range, the accuracies for the spectral slope were consistently higher than for Lempel-Ziv complexity (cf., Figure 4).

Collectively, the results so far suggest that spectral slope and Lempel-Ziv complexity are both sensitive markers that can track brain states during sleep and wakefulness due to changes in sleep depth or due to general task engagement and differences in required cognitive resources. However, while the two parameters are modulated in remarkably similar ways when using a broadband frequency range (1 – 45Hz), they express diverging patterns when a restricted frequency range (30 – 45Hz) is used. Therefore, we next explored the relationship between spectral slope and Lempel-Ziv complexity.

### Relationship between the spectral slope and Lempel-Ziv complexity

First, we assessed the robustness of the spectral slope and Lempel-Ziv complexity estimations across different recordings per subject. We correlated each parameter (in the narrow-and broadband frequency range) with itself between the different lab-sessions for each sleep stage and task. Between all lab-sessions, the parameters were strongly positively correlated, indicating a substantial overlap of information over the different recordings (see Supplementary file-Table 4). To identify the relationship between the spectral slope and Lempel-Ziv complexity for each of the two frequency ranges, we further computed the correlations between the two parameters. In the broadband frequency range, the slope and complexity were consistently positively correlated across all sleep stages and tasks (see Figure 7A and B, right columns). However, this relationship vanished in the narrowband frequency range where the correlation between slope and complexity was inconsistent and ranged from significant negative to positive values (see Figure 7A and B, left columns). These results imply that the two parameters are indeed not very codependent in the narrowband range. In contrast, the information is almost entirely redundant in the broadband frequency range. This fits well to our previous results (cf., Figures 2, 3, 5 and 6) where only the narrowband slope and complexity were differentially modulated by sleep stage and the different tasks. Taken together, this suggests that the narrowband spectral slope and Lempel-Ziv complexity actually track different features of brain activity that are only explicitly captured when using a restricted frequency range as for instance 30 – 45Hz. In broader frequency ranges, the dominance of other, especially lower frequencies might blur these effects.

**Figure 7.**
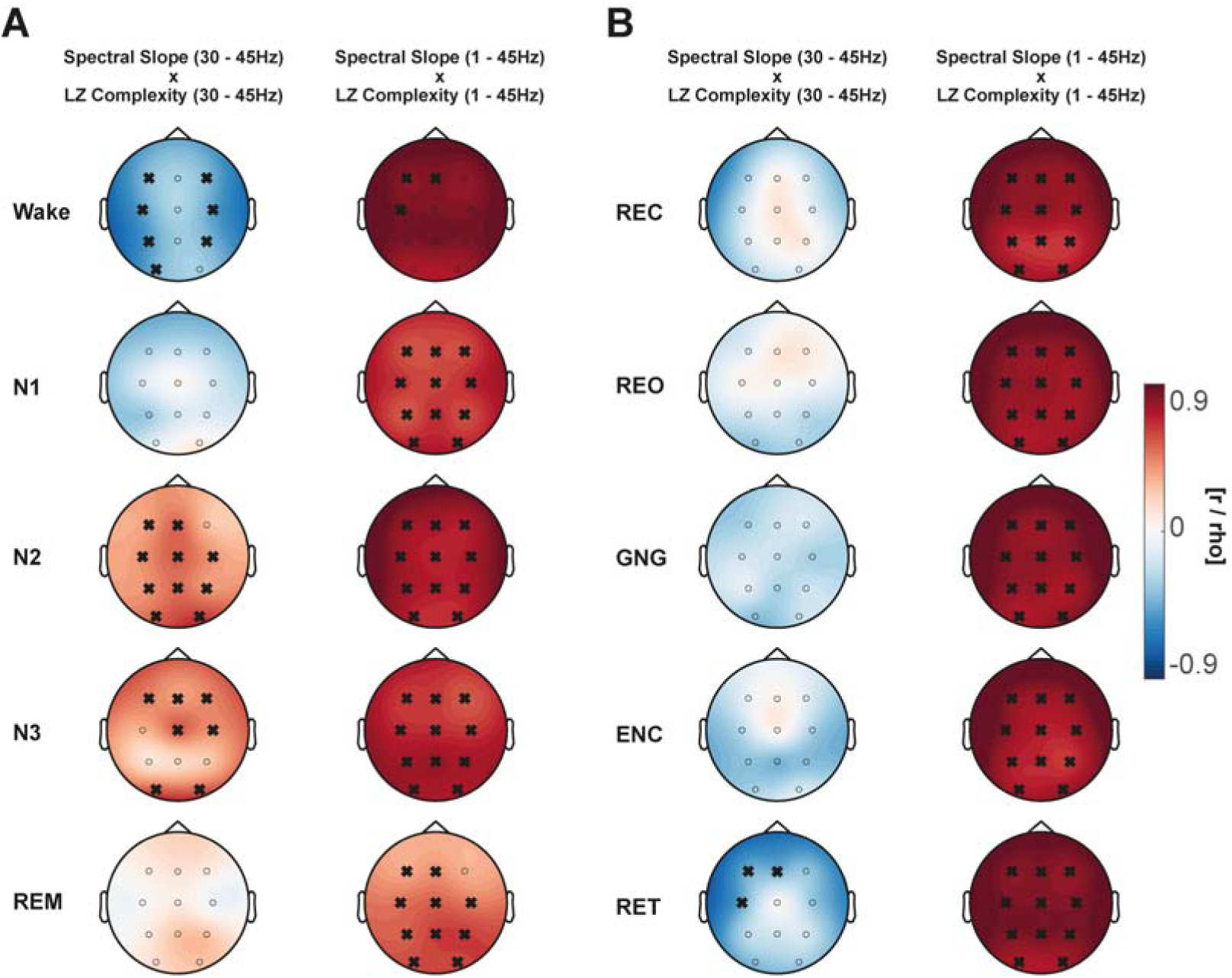
Summary of correlations between the spectral slope and Lempel-Ziv (LZ) complexity from 30 – 45Hz and 1 – 45Hz. The sleep (A) and task (B) data per subject were averaged across all lab-sessions. For task data, only the evening assessments highlighted by the dashed dark-green rectangle in Figure 1 were considered. Significant correlations (*p* ≤.050 after correcting for false discovery rate) are highlighted with a cross on the topographical maps (color codes for the size and directionality of the correlation coefficients). The 30 – 45Hz slope and complexity showed no consistent positive or negative relationship across tasks and sleep stages. In contrast, the 1 – 45Hz slope and complexity were consistently positively correlated over all tasks and sleep stages (*N* = 28). **Figure 7: Supplement 1.** Correlation of the slope and complexity with themselves in the narrow-or broadband frequency range during sleep (A) and wakefulness (B). Only the spectral slope was consistently positively correlated with itself, whereas LZ complexity was slightly negatively correlated with itself between the two frequency ranges.

Finally, we assessed how strongly the spectral slope and Lempel-Ziv complexity were correlated with themselves between the narrow-and broadband frequency range. The narrow-and broadband slopes were consistently positively correlated, whereas the opposite was true for complexity (see Figure 7 – Figure Supplement 1). Thus, flatter narrowband slopes were usually also associated with flatter slopes in the broadband range, but lower narrowband complexity was often even associated with higher broadband complexity.

### The spectral slope as an electrophysiological marker of task performance

Having established that spectral slope and Lempel-Ziv complexity are not only modulated by sleep but also differ between tasks in a frequency range specific manner, we next investigated their relationship with task performance. Thus, we correlated the spectral slope and Lempel-Ziv complexity from the narrow-and broadband frequency range during the Go/Nogo task with the according performance score (percentage of correct trials divided by median reaction time) over multiple sessions. Again, this allowed us to test the robustness of any correlations with behavior. Only flatter slopes in the narrowband range (30 – 45Hz) were consistently related to better task performance (see Figure 8).

**Figure 8.**
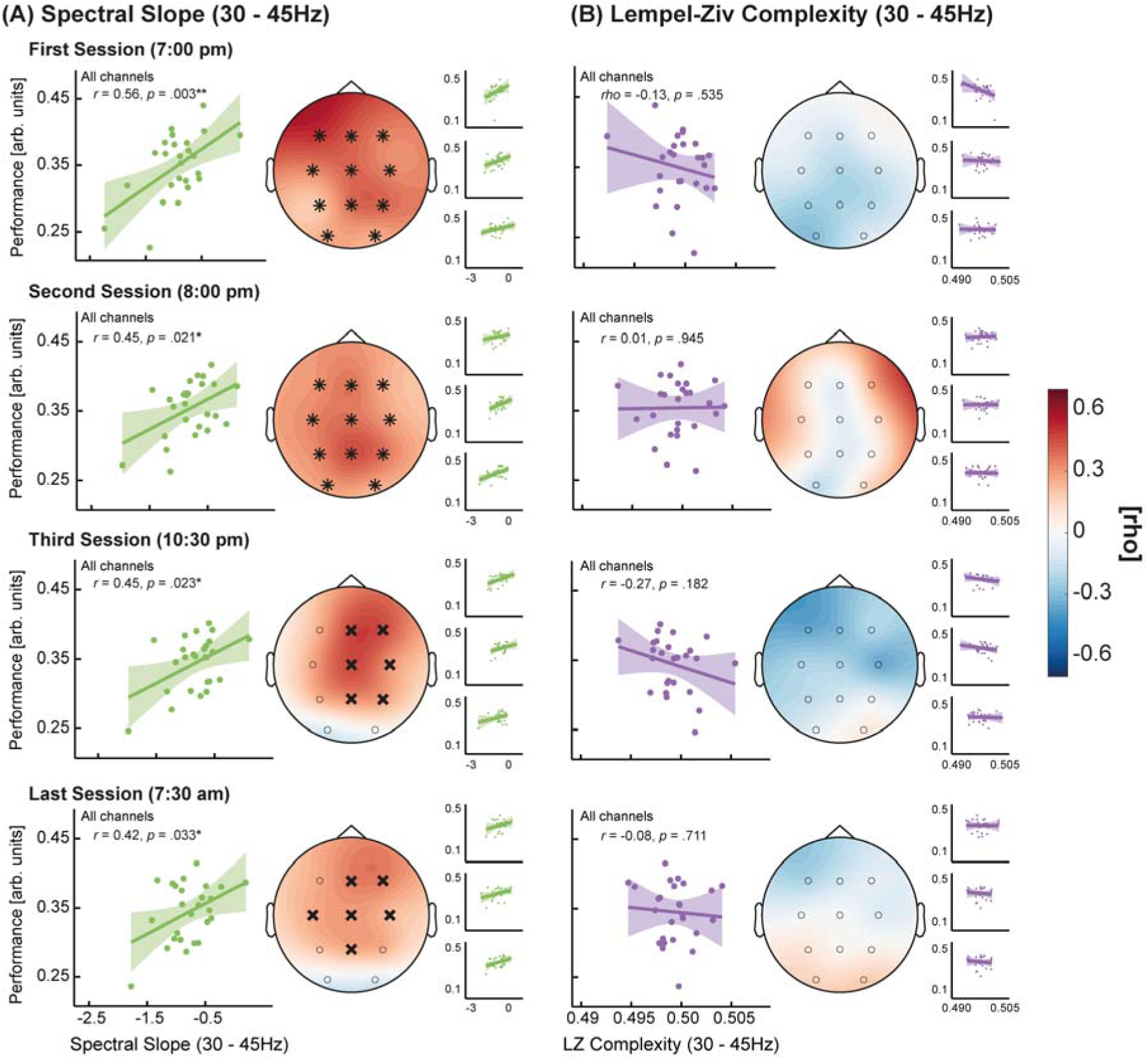
Relationship between Go/Nogo task performance and spectral slope (A) or Lempel-Ziv (LZ) complexity (B) within 30 – 45Hz across different assessment times. For the large scatterplots, data was averaged across all lab-sessions (small scatterplots show the relationship per lab-session). The topoplots depict the correlation strength for each electrode. Electrodes forming a significant cluster are highlighted with asterisks. Those showing a significant correlation after false discovery rate correction but did not from a cluster are marked with a cross. Only the narrowband spectral slope showed a consistent positive relationship with task performance (*N* = 26). **Figure 8: Supplement 1.** Results when using the broadband (1 – 45Hz) frequency range. No significant relationships emerged for the spectral slope and Lempel-Ziv complexity, even though correlations were consistently positive for both parameters.

Lempel-Ziv complexity, on the other hand, did not correlate with performance, neither in the narrow-nor in the broadband (1 – 45Hz) range (see Figure 8 and Figure 8 – Figure Supplement 1). In the broadband range, the relationship with task performance was still consistently positive for both parameters but did not reach statistical significance. The fact that this positive relationship was strengthened and turned significant only for the slope in the narrowband range again suggests a distinct role of the narrowband slope, which might also be interpreted as a specific marker of task performance.

Next, we determined whether the correlation between the narrowband slope and task performance also holds for memory performance in the declarative learning task. Therefore, we correlated the spectral slope and Lempel-Ziv complexity during the retrieval sessions of our declarative memory task with the according recall performance (i.e., percentage of correctly recalled word pairs). Even though the overall pattern was similar to the Go/Nogo task, most correlation coefficients only showed a trend towards statistical significance (see Figure 9).

**Figure 9.**
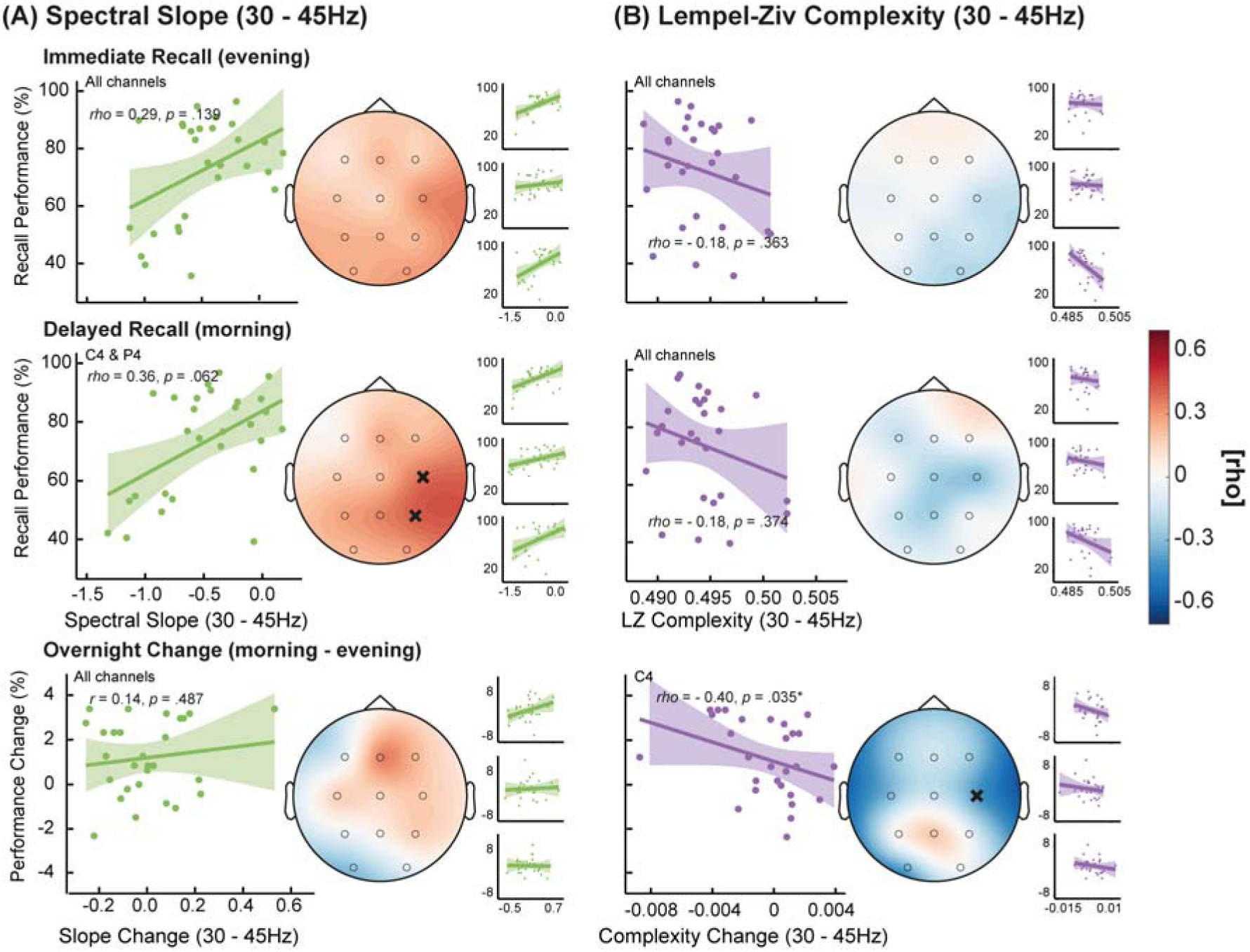
Relationship between declarative memory recall performance and spectral slope (A) or Lempel-Ziv (LZ) complexity (B) within 30 – 45Hz. Results are shown for the immediate recall during the evening and the delayed recall in the next morning as well as for the overnight change. For the large scatterplots, data was averaged across all lab-sessions (small scatterplots show the relationship per session). The topoplots represent the strength of the correlations on each electrode. Even though the spectral slope was consistently positively correlated with recall performance, no electrodes formed a significant cluster. Significant single electrodes that survived false discovery rate correction are highlighted with a cross (*N* = 28). **Figure 9: Supplement 1.** Results when using the broadband 1 – 45Hz frequency range. No relationship observable between recall performance and slope or complexity.

Despite the lack of statistical significance on most electrodes, only the narrowband spectral slope was again consistently positively correlated with recall performance. This indicates that flatter slopes, especially in the narrowband frequency range, are not only related to better attentional performance but might also benefit declarative memory. In contrast, the narrowband complexity was not positively correlated with memory performance and even expressed a negative relationship on some electrodes. Since we observed a positive relationship between overnight decreases in resting state slopes and memory performance in another study (Lendner et al., 2022), we further assessed whether the overnight change in slope during the retrieval task is also correlated with sleep-dependent memory consolidation. However, we did not find a significant relationship, indicating that while flatter slopes during the retrieval were associated with slightly better memory performance, overnight changes in the slope or complexity were not related to performance changes in our study.

In the broadband frequency range, both parameters did not show a consistent relationship with recall performance (see Figure 9 – Figure Supplement 1). Finally, we analyzed whether the similar results between the Go/Nogo and declarative memory task performance could be traced back to better overall attention and higher task engagement but there was no significant relationship between the performance scores from the two tasks (evening: rho = 0.10, *p* =.611; morning: rho = 0.06, *p* =.766). Thus, subjects that performed well in the Go/Nogo task did not necessarily achieve a high recall performance score in the declarative memory task.

## Discussion

In this study comprising three experimental recordings with multiple measurements per subject, we demonstrated that the spectral slope and Lempel-Ziv complexity both reliably delineate sleep stages and are modulated by different cognitive tasks during wakefulness. Critically, we provided evidence that the correlation between spectral slope and Lempel-Ziv complexity strongly depends on the frequency content, which alters their modulation across tasks and sleep stages. The narrowband (30 – 45Hz) spectral slope was best suited to differentiate REM sleep from wakefulness, even though the broadband (1 – 45Hz) slope and complexity were more strongly modulated by sleep stages in general. During wakefulness, active task engagement was associated with flatter slopes in the narrow-and broadband range, but only with higher complexity in the broadband range. In general, the broadband range was also better suited for both parameters to capture the differences between tasks in the classification analyses. Critically, solely the narrowband spectral slope tracked task performance in an auditory attention task (Go/Nogo) as well as in a declarative memory task.

### Sleep stage specific alterations of spectral slope and Lempel-Ziv complexity

Our findings corroborate previous research which demonstrated that the spectral slope and Lempel-Ziv complexity are sensitive markers of sleep depth (Abásolo et al., 2015; Bódizs et al., 2021; Kozhemiako et al., 2022; Lendner et al., 2020; Pascovich et al., 2022; Schartner et al., 2017). Building upon these findings, we leveraged repeated EEG recordings per subject and confirmed that the two parameters can robustly differentiate all sleep stages from wakefulness. Overall, sleep stages could be better delineated when a broadband frequency range (1 – 45Hz) was used for calculation of the spectral slope and Lempel-Ziv complexity. This is probably due to the fact that the broadband range encompasses the frequencies typically used for traditional sleep scoring, such as slow wave activity (0.5 – 4Hz) and sleep spindles (11 – 15Hz; Dijk, 1995), thereby increasing the sleep stage specific information in the underlying signal. However, only the spectral slope within the narrowband frequency range (30 – 45Hz) clearly distinguished REM sleep from all other sleep stages, which is in line with recent findings by Lendner et al. (2020). This behavior of the narrowband spectral slope contradicted the overall modulation of slope and complexity in the broadband range, where both parameters showed a relative, more wake-like, increase during REM sleep. Since REM sleep (sometimes called ‘paradoxical sleep’; Peigneux et al., 2001 or Siegel, 2011) is in comparison to NREM sleep characterized by less prominent oscillations and a more desynchronized EEG pattern similar to wakefulness (Blumberg et al., 2020; Peever & Fuller, 2017), these disparate results between the two frequency ranges suggest that the narrowband slope mainly measures non-oscillatory, aperiodic brain activity. The relative increase in broadband complexity during REM sleep has been attributed to higher levels of conscious content that accompany vivid dreaming and thus require more complex brain activity than deeper, mostly dreamless sleep stages (Lau et al., 2022; Mateos et al., 2018).

Recent modeling work has also linked especially the narrowband spectral slope with the excitation to inhibition (E/I) balance in the brain (Gao et al., 2017). Within this framework, steeper slopes during REM sleep potentially reflect stronger inhibitory brain activity. This might allow the brain to decouple from its environment and, by maintaining muscle atonia, to enable the consolidation of emotional memories and the experience of vivid dreams (Aime et al., 2022) without the danger of acting them out. The narrowband (30 – 45Hz) complexity, however, expressed a diverging pattern compared to the narrowband slope and stayed almost constant across all sleep stages with even a slight increase from N1 to N2 sleep. Even though this is one of the first studies that compared the spectral slope and Lempel-Ziv complexity during sleep, the congruency of both measures within the broadband frequency range might not be surprising, since other studies investigating the two parameters individually have shown their decrease across sleep (Aamodt et al., 2021; Lendner et al., 2020; Miskovic et al., 2019; Pereda et al., 1998; Schartner et al., 2017).

Although we were able to classify sleep stages consistently above chance level with both parameters, it should be noted that our classifier was trained and tested only on our own data. Furthermore, we did not compare the performance of the spectral slope and Lempel-Ziv complexity to other potentially powerful biomarkers (e.g., heart rate variability or blood pressure as utilized by Kuula & Pesonen, 2021; Mitsukura et al., 2020; Radha et al., 2019 or van de Borne et al., 1994). Therefore, it would be interesting to see how accurately sleep stages can be scored exclusively by means of the slope or complexity and how these two markers perform in comparison to other indices of sleep depth or accelerometric data from actigraphy (Lüdtke et al., 2021; Sadeh et al., 1989) and multisensor consumer-wearables (Ameen et al., 2019; Boe et al., 2019; Roberts et al., 2020; Tal et al., 2017).

### Spectral slope and Lempel-Ziv complexity are modulated by task engagement

In addition to our findings during sleep, we demonstrate that the spectral slope and Lempel-Ziv complexity can track different cognitive tasks and are affected by task engagement in general. That slope and complexity are generally also modulated during wakefulness is in line with other research (Jacob et al., 2021; Mediano et al., 2021; Sheehan et al., 2018; Waschke et al., 2021), however, to our best knowledge this is the first study assessing multiple cognitive tasks and different resting conditions as well as the influence of different frequency ranges on the discrimination accuracy of the two parameters. Similar to sleep, we observed a homogenous modulation of the broadband (1 – 45Hz) slope and complexity, where flatter slopes and higher complexity were associated with active task engagement compared to resting. In addition, especially the retrieval session of the declarative learning task appeared to yield the flattest broadband slope and highest complexity in comparison to the encoding session or an auditory Go/Nogo task. This pattern was identical for the narrowband slope but was inverted for the narrowband complexity. In the E/I balance framework, flatter slopes are the result of higher excitation in the brain (Chini et al., 2022; Gao et al., 2017). Thus, our observed pattern of a flattening of the spectral slope with task engagement and between different tasks might suggest that this effect could maybe be attributed to differences in the amount of required cognitive resources for the different tasks which might lead to stronger excitatory brain activity (Harris & Thiele, 2011; He, 2011; Kanashiro et al., 2017). Unlike Waschke et al. (2021), who reported a stronger occipital flattening of the slope in a visual compared to an auditory task, we did not observe clear topographical differences between modalities, even though the attentional Go/Nogo task was entirely auditory except for a fixation-cross whereas the declarative memory task mainly relied on visual content. However, this lack of topographical distinctiveness might also be due to a partial overlap between involved brain areas since both, auditory discrimination and learning involve frontotemporal brain regions (Ackerman, 1992; Halsband, 1998).

### Differential contributions of narrow-and broadband frequency ranges

Based on the results from the broadband frequency range, it is tempting to assume that the spectral slope and Lempel-Ziv complexity are indexing the same or at least very similar features of brain activity. Indeed, according to Medel et al. (2020), both parameters might actually be driven by the transition entropy of the underlying cortical system and flatter slopes as well as higher complexity values could be similarly characteristic of the same cortical states. However, the divergence between the narrow-and broadband slope and complexity during sleep and wakefulness clearly demonstrates that the two parameters cannot be used interchangeable. Instead, especially in a restricted frequency range, they track different facets of the underlying brain activity. Here, we revealed that the selected frequency range dramatically influences the information that the two parameters provide and therefore also their interrelation. Using a narrowband frequency range from 30 – 45Hz for estimation decreases the relationship between the spectral slope and Lempel-Ziv complexity. During wakefulness, different contributions of oscillatory and aperiodic brain activity might account for their diverging patterns in the narrowband range. At first, it appears paradoxical that flatter narrowband slopes, representing an increase in aperiodic activity, should be accompanied by a decrease in complexity since complexity should also increase with higher signal irregularity. However, others have also reported this type of counterintuitive behavior of Lempel-Ziv complexity. Mediano et al. (2021) showed that in MEG within 0.5 – 30Hz, active tasks actually exhibited lower complexity values compared to rested wakefulness. Additionally, a recent review from Lau et al. (2022) discussed several studies that reported apparently contradicting modulations of signal complexity in different clinical conditions, where some report lower and others higher levels of complexity. Thus, the question whether higher signal complexity can always be clearly interpreted as more complex or irregular brain activity remains unclear. So far, the best explanation for the contradictory findings in the complexity literature is that higher complexity values can both represent either more complex or more random systems (La Torre-Luque et al., 2016), which makes it difficult to argue whether higher complexity always represents a healthier brain. Interestingly, other studies also showed a strong relationship between different complexity or entropy measures and the spectral slope (Colombo et al., 2019; Miskovic et al., 2019; Waschke et al., 2017), thus, it would be interesting to investigate in the future what drives their shared information and under which circumstances (i.e., frequency ranges) this relationship vanishes.

### The narrowband spectral slope as a unique marker of task performance

When relating the spectral slope and Lempel-Ziv complexity to behavioral outcomes, we observed that only the narrowband slope within 30 – 45Hz was correlated consistently with attentional task performance in an auditory Go/Nogo task across all recordings per subject. Thus, it appears that the narrowband slope serves as a particularly sensitive marker for task-dependent fluctuations in brain states associated with behavioral performance. This association between adaptively flatter slopes and better task performance might even translate to more general cognitive tasks that do not solely rely on attention since we also observed a consistent positive but weaker relationship with memory performance. In larger scale studies that rely on databases or in multicenter studies, which commonly have higher statistical power, however, the broadband slope and complexity were also significantly correlated with task performance. For instance, Mediano et al. (2021) and Waschke et al. (2021) found an association between task-specific attention levels and spectral slope or Lempel-Ziv complexity in a broader frequency range. As in our study the correlation between the broadband slope and complexity with the Go/Nogo task performance was also consistently positive but too weak to reach statistical significance, these findings do not necessarily contradict our claim that the narrowband spectral slope is even more sensitive to adaptive task-dependent changes in brain state. In contrast, this shows that lower statistical power might suffice for the narrowband slope to index robust relationships with behavioral performance.

### Limitations

It should be noted that the cognitive tasks in this work were not specifically designed for the analysis of varying levels of task demands or difficulty as the data presented here was obtained from a study that was originally designed for the investigation of short-wavelength light effects on sleep, attention and memory performance (cf., Höhn et al., 2021 and Schmid et al., 2021). Even though it is tempting to attribute the task differences to variations in task demand as the participants reported differences in task difficulty that were not systematically measured, this would overlook other significant factors such as differences in task modality. This therefore limits the conclusion that the neural differences between the Go/Nogo and learning tasks can be ascribed to the level of task demand. While there is evidence in the literature that attentional tasks and learning tasks do differ in their level of cognitive demand (Bambrah et al., 2019; Sweller, 2011), it also seems to be highly dependent on the specific task instructions and modalities Thus, it would be necessary to systematically assess the subjective levels of task demand and difficulty for each task used in the present study to be able to ascribe task differences to specific task characteristics. In the future, it might also be promising to contrast tasks that exclusively rely on different cognitive resources and sensory modalities (e.g., auditory vs. visual) to assess how spectral slope and Lempel-Ziv complexity adapt topographically to different modalities.

Even though we used only 11 scalp electrodes, we still robustly detected the effects of sleep stage and task demand, providing evidence for the power of the slope and complexity as indices of different brain states. Nevertheless, research with high-density or intracranial EEG might further contribute to the understanding of which topographical areas are most influential in driving changes in slope or complexity across brain states.

Finally, we only recruited healthy biologically male adults in a restricted age range (18 – 25 years) in order to avoid potential sex differences and hormonal effects (Kozhemiako et al., 2022; Plamberger et al., 2021). Therefore, it is unclear to what extent our results generalize to other populations. While sex does not necessarily affect the slope or complexity when controlling for overall signal amplitude (Bódizs et al., 2021; Tosun et al., 2019), age does seem to play an important role in terms of developmental changes in the spectral slope and decorrelation of brain activity, which begins during early childhood (Chini et al., 2022; Schaworonkow & Voytek, 2021) and lasts until late adulthood (Dave et al., 2018). While this task-independent flattening of the slope in older subjects has been associated with decline in cognitive functioning (Voytek et al., 2015), our results suggest that task-dependent increases in excitation (expressed by flatter slopes) might be beneficial for behavioral performance. Thus, an adaptive task-specific modulation of the slope in healthy individuals appears to be associated with better task performance and might index cognitive adaptability.

## Conclusions

Taken together, our results demonstrate that the EEG spectral slope and Lempel-Ziv complexity are powerful indices of different brain states during sleep and wakefulness. We provide robust evidence from multiple recordings of three within-subjects measurements, showing that sleep stages and different cognitive tasks are reliably indexed by both, the spectral slope and Lempel-Ziv complexity. Critically, we show that the selected frequency range has a strong impact on the interpretability and functional relevance of the two parameters. During sleep and wakefulness, the broadband slope and complexity are more sensitive and better suited to distinguish between different brain states across vigilance states and tasks. However, only the narrowband spectral slope within 30 – 45Hz turned out to be a powerful index of behavioral performance and was best suited to differentiate REM sleep from wakefulness and all other sleep stages.

## Materials and methods

**Key resources table.**
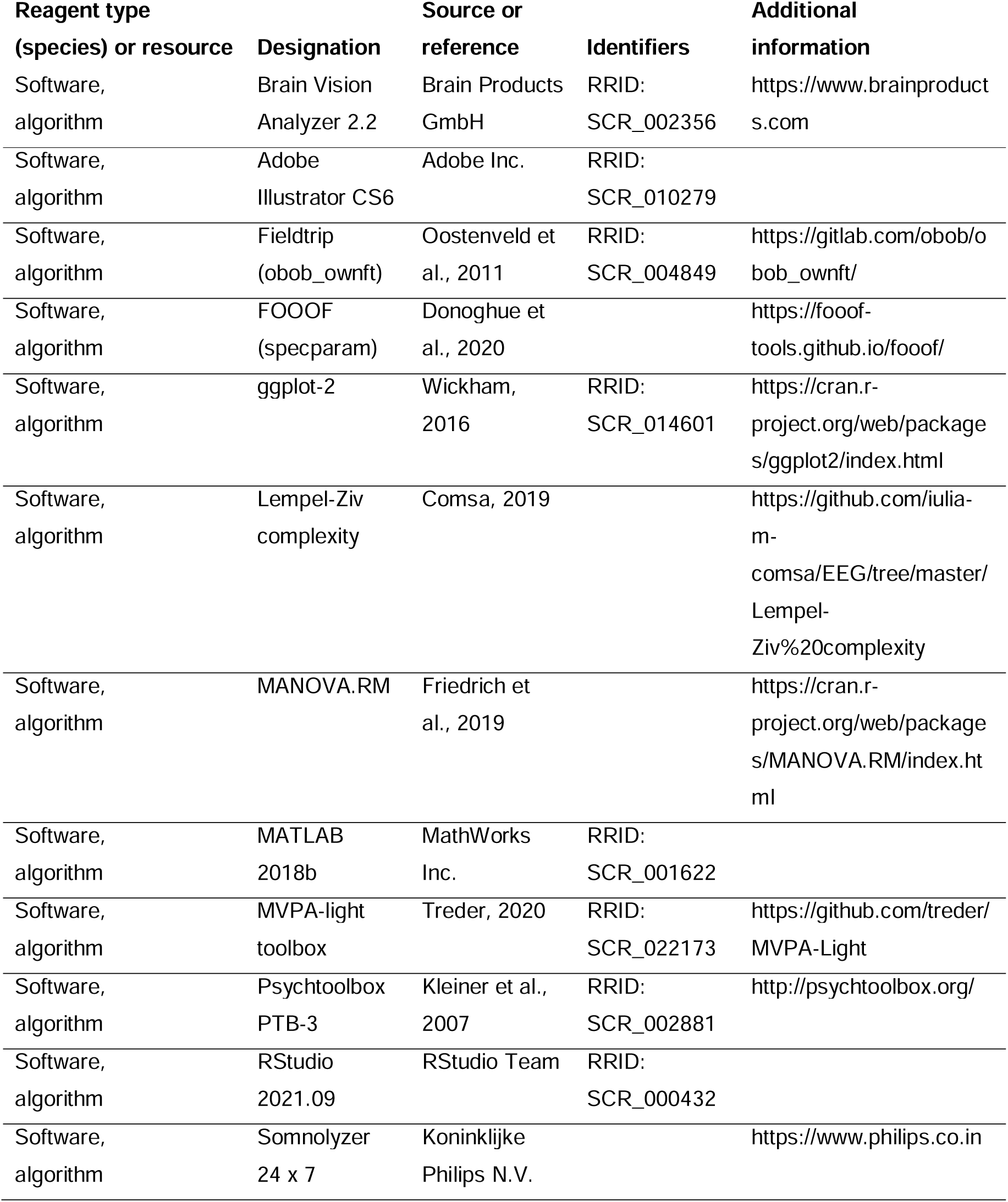

### Participants and inclusion criteria

We recorded data from 28 biologically male participants (18 – 25 years; mean age 21.54 ± 1.90 years). Final sample sizes varied for each analysis between N = 26 – 28 as some participants had missing data for specific tasks or timepoints (the exact sample size for each analysis is provided in the corresponding figure captions). All participants were free of medication and did not suffer from a mental or physiological illness or from sleep problems. They adhered to a regular sleep-wake rhythm (i.e., regular bedtimes with about 8 hours of sleep per night) and refrained from drug abuse and above-average caffeine consumption (more than three cups of coffee per day) during participation. For screening purposes, all subjects filled in an entrance questionnaire in which we checked for sleep quality, mood, anxiety, perceived stress level and chronotype (see Supplementary file – Table 1). Written informed consent was obtained from every participant and all subjects were remunerated with either 100€ and 16 hours course credit or 50€ and 24 hours course credit. The study was approved by the local ethics committee of the University of Salzburg (EK-GZ: 16/2014) and conducted in agreement with the Declaration of Helsinki.

### Experimental protocol

#### Study design

Each subject participated over a time span of 14 days, with an entrance examination marking day one (an outline of the study protocol is presented in Figure 1). From that day on, participants wore an actigraph (MotionWatch 8; CamNtech Ltd, Cambridge, England) and filled in daily online sleep protocols (LimeSurvey GmbH, Hamburg, Germany), which we used to check for compliance with a regular sleep-wake rhythm.

The first recording was scheduled on day four and was implemented only for adaptation purposes in order to avoid potential first night effects (Browman & Cartwright, 1980; Curcio et al., 2004). After placement of all EEG, ECG, EMG and EOG electrodes, the participants were familiarized with the resting and Go/Nogo tasks. Bedtime was scheduled for approximately 11:00 pm and the participants were woken up 8 hours after lights out before they left the laboratory at approximately 9:00 am.

The experimental recordings were scheduled on days 7, 10 and 13. Participants arrived at 6:00 pm and EEG, ECG, EMG and EOG electrodes were mounted. The recordings started with an initial resting session (3min eyes closed and 3min eyes open) and the Go/Nogo task (10min), which was followed by the encoding sessions (two times 14min) of a declarative memory task. Before the first cued recall, another resting and Go/Nogo session were conducted. Afterwards, the participants had a 1.5 hours break from the tasks, in which they read stories under different light conditions (for details cf., Schmid et al., 2021). Before going to bed at approximately 11:00 pm, participants completed the last resting and Go/Nogo session of the day. After awakening, a morning session of resting and the Go/Nogo task as well as another recall from the declarative memory task were performed. During all wake-recordings, daylight mimicking room lights (provided by Emilum GmbH, Oberalm, Austria) were dimmed to 4.5 photopic lux and room temperature was adjusted via air conditioning based on participant’s preferences.

#### Go/Nogo task

To assess objective levels of attention, we implemented an auditory version of the Go/Nogo paradigm (Donders, 1969) via the Psychophysics Toolbox (PTB-3; Kleiner et al., 2007) in MATLAB (Release 2018b, The MathWorks Inc., Natick, MA). Participants were asked to react as quickly as possible with a button press on a response time box (RTBox v5/6; Ohio State University, Columbus, OH) whenever they heard a ‘Go’ sound and needed to inhibit their reaction when a ‘Nogo’ sound was played. The task comprised 400 trials with Go sounds being presented in 80% of the trials and Nogo sounds occurring in the remaining 20% of trials (the order of Go and Nogo sounds was randomized each time). The two stimuli used for the Go and Nogo sounds were low-(1000Hz) and high-pitched (1500Hz) tones, which were presented for 50ms with a varying interstimulus interval (1480 – 1880ms). Whether the low-or high-pitched sound represented the Go-signal was determined by chance at the beginning of each session. Participants had to react within 500ms for the response to be considered valid, but responses were recorded until 1000ms post-stimulus with reaction times longer than 500ms being regarded as attentional lapses. From each session, the performance score was computed by dividing the percentage of correct trials by the median reaction time of all valid responses (≤ 500ms, no errors) in milliseconds (Figueiro et al., 2016; Höhn et al., 2021).

#### Declarative memory task

Participants encoded a set of 80 word pairs on days 7, 10 and 13. To avoid learning effects over time, a different but similarly difficult set of 80 word pairs was presented on each of the three days. The order of the sets was randomized across subjects. Each set was presented twice for 14min during encoding and the data from both encoding sessions was pooled for further analyses. Each word pair was presented for 1500ms and was followed by a fixation-cross for 8500ms. Participants were instructed to encode the word pair as vividly as possible during the presentation of the fixation-cross by imagining a semantic connection between the two words. During the cued recall sessions, only the first word of a pair was presented, and participants were asked to press a button on the response time box as soon as they remembered the second word. Whenever a button was pressed, the participant was instructed to name the missing word and a fixation-cross appeared for 3500ms while the experimenter noted the answer. When no button was pressed, the fixation-cross appeared automatically after 6500ms. Recall performance was measured as the percentage of correctly recalled word pairs during each retrieval session. To assess the overnight change in recall performance, the increase in percentage from the performance in the evening to the performance the following morning was computed.

### EEG recording and analyses

All electrophysiological data were recorded with a sampling rate of 500Hz via the BrainVision Recorder software (Version 2.11, Brain Products GmbH, 2015) using a 32 channel BrainAmp system (Brain Products GmbH, Munich, Germany). We placed 11 gold-cup electrodes (Grass Technologies, Astro-Med GmbH, Rodgau, Germany) according to the international 10-20 system on the positions: F3, Fz, F4, C3, Cz, C4, P3, Pz, P4, O1 and O2. The average of positions A1 and A2 on the left and right mastoids was used for offline re-referencing as the data were online referenced against Cz. Fpz was used as ground electrode. Additionally, two EMG electrodes were placed on the musculus mentalis for measuring muscle activity during sleep and four EOG electrodes around the eyes to record horizontal and vertical eye movements. ECG was recorded with an electrode on the right clavicular and another one on the lowest left costal arch. Impedances were always kept below 10kΩ.

#### Polysomnography

The time in bed was standardized for all polysomnography recordings and comprised 8 hours. For sleep staging, the data were first low-pass filtered at 30Hz and re-referenced to contralateral mastoids with the BrainVision Analyzer software (Version 2.2.0.7383, Brain Products GmbH, 2019). Physio-channels were referenced in a bipolar manner and the data were down-sampled to 128Hz before sleep stages were classified for each 30 second epoch with the Somnolyzer 24 x 7 algorithm (Koninklijke Philips N.V.; Eindhoven, The Netherlands) in accordance with the criteria of the American Academy of Sleep Medicine (Richard et al., 2012). The results were finally verified by a human expert scorer. The general sleep architecture of each night is presented descriptively in Supplementary file – Table 5.

#### EEG preprocessing

In a first step, the data were processed with the BrainVision Analyzer software, and we applied a 0.3Hz high-pass as well as a 50Hz notch filter. The EEG channels were re-referenced to linked mastoids and the online reference Cz was restored. We corrected for eye movements with the Gratton & Coles method (Gratton et al., 1983; only implemented for data during wakefulness) and ran an automatic artifact detection procedure on all scalp EEG channels, which was manually checked afterwards. Events with a voltage step exceeding 50μV/ms, an absolute voltage difference of more than 400μV within 200ms or activity lower than 0.5μV for at least 100ms were automatically marked as bad intervals. The voltage difference criterion is slightly more lenient than the default recommendation from the Brain Vision Analyzer (Brain Products GmbH, 2019) to avoid false positive classifications during slow wave sleep. If severe muscle or movement artefacts were missed by the automatic detection, they were also marked manually. The data were then down-sampled to 250Hz and exported for further analyses in MATLAB. The continuous data were subsequently segmented into epochs of 4s for each task and sleep stage using the fieldtrip toolbox (Oostenveld et al., 2011). To be able to compare all task-and sleep-data, we decided to set the epoch-length to 4s as this enabled the best tradeoff between sufficient epochs even for the shortest tasks (3min resting sessions) and an adequate frequency resolution within 0.5 – 45Hz. All artifact-containing epochs (defined as > 1% being detected as artifact) were removed for the following analyses. Since the remaining number of clean epochs from the different tasks (resting, Go/Nogo, encoding and retrieval) and sleep-stages (WAKE, N1, N2, N3 and REM) varied dramatically due to different recording lengths, we balanced the number of epochs across tasks and sleep-stages for the multivariate pattern analyses (MVPA) to ensure the validity of the classification results. In more detail, we set the maximum number of epochs for the MVPA analyses to the highest possible number of epochs from the shortest task (i.e., 45 epochs as the resting sessions only comprised 3min). To do so, we drew a random subset of 45 epochs from all data that contained more than 45 clean epochs. For all other analyses we used all available data to maximize the signal to noise ratio wherever possible (for the number of epochs used per task and sleep stage see Supplementary file – Table 6).

#### Spectral Slope

To obtain the spectral slope, we first calculated power-spectra between 0.5 – 45Hz from the preprocessed, 4s segmented data via the mtmfft method in Fieldtrip (Oostenveld et al., 2011) using a multi-taper approach (1Hz frequency smoothing; Lendner et al., 2020). To extract the spectral slope information, we applied robust linear fits (using the robust fit MATLAB function) in log-log space between 30 – 45Hz based on a previously established method (Lendner et al., 2020). We decided to use robust linear fits instead of using the FOOOF algorithm (alternatively known as specparam; Donoghue et al., 2020) for the narrowband frequency range since this approach has already been established to yield a sensitive aperiodic marker of arousal by Lendner et al. (2020) and because in this frequency range also the FOOOF would approximate a linear fit, thus leading to highly comparable results. However, for the broadband frequency range (1 – 45Hz), we applied the FOOOF algorithm to extract the slope since linear fits would have been skewed significantly by oscillatory bumps in the power spectrum.

#### Lempel-Ziv Complexity

We followed previous approaches (Mediano et al., 2021; Schartner et al., 2015) and calculated the Lempel-Ziv-Welch complexity (Lempel & Ziv, 1976; Welch, 1984) as a proxy for signal complexity per channel and epoch. To obtain the complexity in the same frequency ranges in which we calculated the spectral slope, we applied additional 1Hz or 30Hz high-pass and 45Hz low-pass filters to ensure that the underlying signal contained the same frequencies as for the spectral slope. As Rivolta et al. (2014) demonstrated that 1000 datapoints are sufficient for reliable Lempel-Ziv complexity analyses during sleep, we used the same 4s segmented data (which translates to 1000 sampling points per epoch in the down sampled data) for the complexity analyses that we used for the spectral slope. We then applied a Hilbert-transformation on each epoch to obtain the instantaneous amplitude. Afterwards, we binarized the resulting single epoch data around its median amplitude and transformed it into a binary sequence. Values of 1 were given for amplitude samples above the median and values of 0 for amplitudes below (or equal with) the median. This binary sequence of ones and zeros was finally subjected to the Lempel-Ziv-Welch complexity algorithm (Comsa, 2019) in MATLAB.

### Statistical analyses

Statistics were calculated in R-Studio (Version 4.1.2.; RStudio Team, 2021). MATLAB functions from the Fieldtrip toolbox and the ggplot-framework (Wickham, 2016) in R were adapted for data visualization.

#### Factorial analyses and correlations

All analyses involved three repeated measurements (on days 7, 10 and 13; cf., Figure 1) and therefore at least two factors (lab-session and task or sleep stage). Since in most cases at least one assumption for parametrical testing was violated, we decided to compute more conservative semi-parametrical analyses with the MANOVA.RM package (Friedrich et al., 2019). For these factorial analyses, data were averaged over all EEG electrodes to facilitate interpretation of the results. In the statistical results, we always refer to the Wald-Type-Statistics (*WTS*) with empirical p-values obtained from permutation resampling procedures and 10.000 iterations. Whenever multiple comparisons were conducted for follow-up testing, p-values were corrected for alpha error inflation with the Benjamini-Hochberg procedure (Benjamini & Hochberg, 1995).

For correlation analyses, we computed Spearman rho coefficients instead of Pearson correlations whenever the normality assumption was significantly violated (indicated by Shapiro-Wilk tests) and in general for all cluster correlations on the whole scalp level. For the cluster corrected correlation approach, we used the Monte-Carlo method with 10.000 iterations to assess the relationship between the EEG parameters per channel and the behavioral measures.

#### Multivariate pattern analyses (MVPA)

Since it is difficult to take topographical patterns into account in classical factorial designs, we additionally computed multivariate pattern analyses using the MVPA-Light toolbox (Treder, 2020) in MATLAB to further exploit the information present in the complexity and slope data as patterns across electrodes. For each task and sleep stage, the complexity and slope from every epoch and electrode was fed into the classifier. Thus, the single epochs per subject were used for training and testing while the complexity and slope patterns over electrodes represented the multivariate information. For comparisons between more than two tasks or sleep stages, multiclass linear-discriminant analyses (LDA) were used and regular LDA for two-condition comparisons. We calculated classifier accuracies per subject via leave-one-out cross validation (LOO-CV) to account for the restricted amount of data available for training and testing in our sample. Since no effects regarding the different lab-sessions emerged, we pooled the data from the different lab-sessions for each subject in order to improve the reliability of the MVPA analyses.

## Acknowledgements

This research was funded by the Austrian Science Fund (FWF, P32028) and the Centre for Cognitive Neuroscience Salzburg (CCNS). C.H. further received funding from the Doctoral College “Imaging the Mind” (FWF; W1233-B). J.D.L. received a grant from the German Research Foundation (DFG LE 3863/2-1). We would like to thank all volunteering participants for their time and effort. Further, we are very grateful for the support from Sarah R. Schmid, Selina Schindlmayr, Daniela Niebler, Lucy Matthews, Marina Thierauf, Leoni Bernstorf, Lorenz Rapp, Henrik Rheinwald and Leonard van Dyck regarding the data collection process and recruitment of participants.

## Additional Information

### Declaration of interest

None.

### Funding

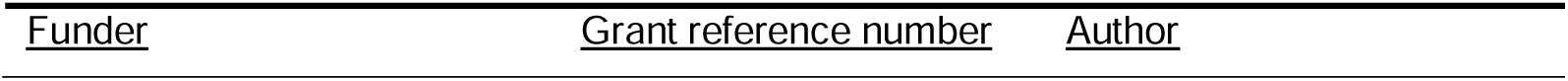
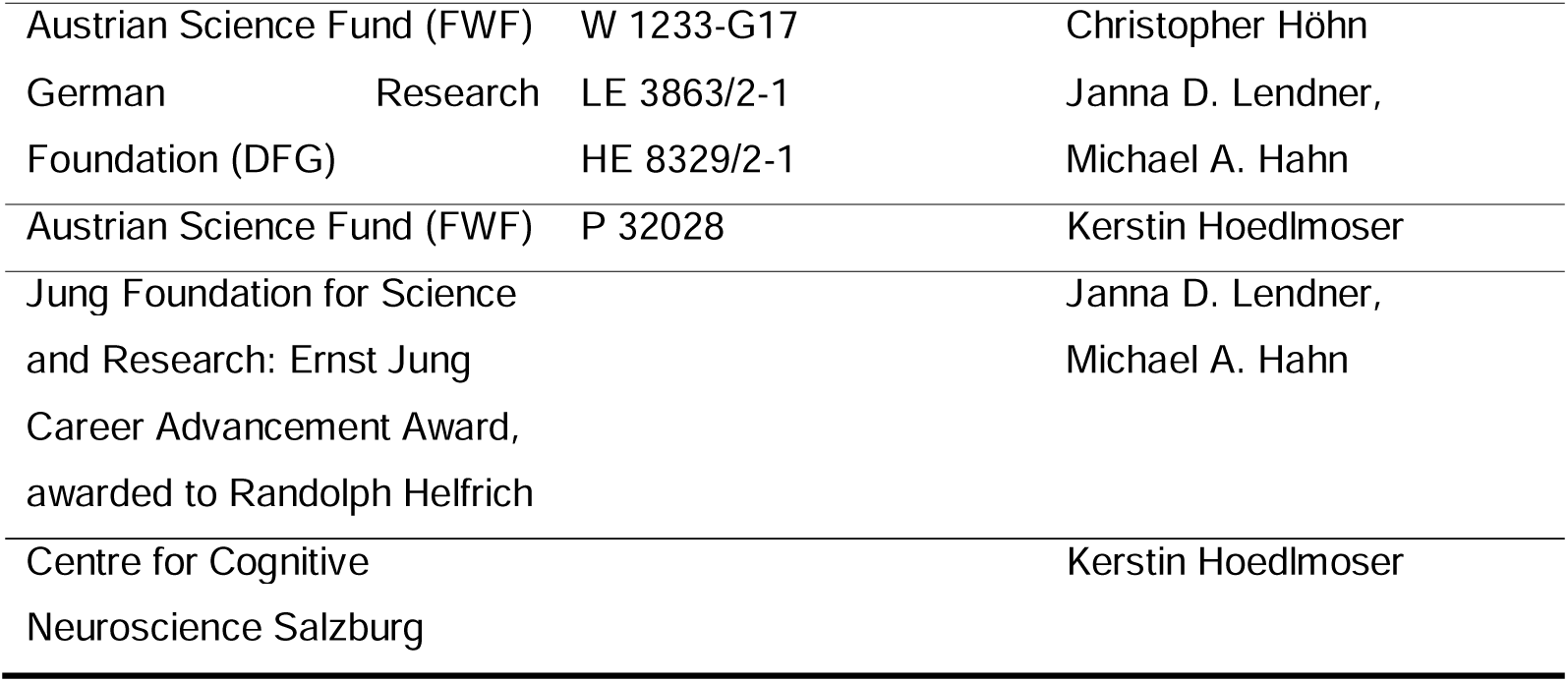

### Ethics

This study was conducted in accordance with the guidelines from the Declaration of Helsinki. Written approval was additionally provided by the local ethics committee of the University of Salzburg (EK-GZ: 16/2014).

## Notes

### Competing Interest Statement

The authors have declared no competing interest.

### Summary of Updates

The manuscript has been updated extensively to incorporate reviewer's comments and suggestions. Additional analyses were added, showing that the modulation of the spectral slope and Lempel-Ziv complexity is not driven by EMG activity. Additionally, the claim of different levels of task-demand has been down tuned as it cannot be shown that the differences in slope and complexity between tasks can be ascribed to varying task demands. Finally, several text passages have been rephrased and additional information is provided in the methods section.

## References

Aamodt, A., Nilsen, A. S., Thürer, B., Moghadam, F. H., Kauppi, N., Juel, B. E., & Storm, J. F. (2021). Eeg Signal Diversity Varies With Sleep Stage and Aspects of Dream Experience. Frontiers in Psychology, 12, 655884. https://doi.org/10.3389/fpsyg.2021.655884

Aarabi, A., & He, B. (2012). A rule-based seizure prediction method for focal neocortical epilepsy. Clinical Neurophysiology, 123(6), 1111–1122. https://doi.org/10.1016/j.clinph.2012.01.014

Abásolo, D., Simons, S., Da Morgado Silva, R., Tononi, G., & Vyazovskiy, V. V. (2015). Lempel-Ziv complexity of cortical activity during sleep and waking in rats. Journal of Neurophysiology, 113(7), 2742–2752. https://doi.org/10.1152/jn.00575.2014

Ackerman, S. (1992). Discovering the Brain. National Academies Press. http://gbv.eblib.com/patron/FullRecord.aspx?p=3376517

Aime, M., Calcini, N., Borsa, M., Campelo, T., Rusterholz, T., Sattin, A., Fellin, T., & Adamantidis, A. (2022). Paradoxical somatodendritic decoupling supports cortical plasticity during REM sleep. Science, 376(6594), 724–730. https://doi.org/10.1126/science.abk2734

Ameen, M. S., Cheung, L. M., Hauser, T., Hahn, M. A., & Schabus, M. (2019). About the Accuracy and Problems of Consumer Devices in the Assessment of Sleep. Sensors, 19(19). https://doi.org/10.3390/s19194160

Andrillon, T., Poulsen, A. T., Hansen, L. K., Léger, D., & Kouider, S. (2016). Neural Markers of Responsiveness to the Environment in Human Sleep. The Journal of Neuroscience, 36(24), 6583–6596. https://doi.org/10.1523/JNEUROSCI.0902-16.2016

Bambrah, V., Hsu, C.l.zF., Toplak, M. E., & Eastwood, J. D. (2019). Anticipated, experienced, and remembered subjective effort and discomfort on sustained attention versus working memory tasks. Consciousness and Cognition, 75, 102812. https://doi.org/10.1016/j.concog.2019.102812

Benjamini, Y., & Hochberg, Y. (1995). Controlling the False Discovery Rate: A Practical and Powerful Approach to Multiple Testing. Journal of the Royal Statistical Society: Series B (Methodological), 57(1), 289–300. https://doi.org/10.1111/j.2517-6161.1995.tb02031.x

Blumberg, M. S., Lesku, J. A., Libourel, P.l.zA., Schmidt, M. H., & Rattenborg, N. C. (2020). What Is REM Sleep? Current Biology, 30(1), R38–R49. https://doi.org/10.1016/j.cub.2019.11.045

Bódizs, R., Szalárdy, O., Horváth, C., Ujma, P. P., Gombos, F., Simor, P., Pótári, A., Zeising, M., Steiger, A., & Dresler, M. (2021). A set of composite, non-redundant EEG measures of NREM sleep based on the power law scaling of the Fourier spectrum. Scientific Reports, 11(1), 2041. https://doi.org/10.1038/s41598-021-81230-7

Boe, A. J., McGee Koch, L. L., O’Brien, M. K., Shawen, N., Rogers, J. A., Lieber, R. L., Reid, K. J., Zee, P. C., & Jayaraman, A. (2019). Automating sleep stage classification using wireless, wearable sensors. NPJ Digital Medicine, 2, 131. https://doi.org/10.1038/s41746-019-0210-1

Brain Products GmbH. (2019). Analyzer | User Manual: Software version 2.2.0 (1st ed.) [PDF].

Browman, C. P., & Cartwright, R. D. (1980). The first-night effect on sleep and dreams. Biological Psychiatry, 15(5), 809–812.

Chini, M., Pfeffer, T., & Hanganu-Opatz, I. (2022). An increase of inhibition drives the developmental decorrelation of neural activity. ELife, 11. https://doi.org/10.7554/eLife.78811

Colombo, M. A., Napolitani, M., Boly, M., Gosseries, O., Casarotto, S., Rosanova, M., Brichant, J.l.zF., Boveroux, P., Rex, S., Laureys, S., Massimini, M., Chieregato, A., & Sarasso, S. (2019). The spectral exponent of the resting EEG indexes the presence of consciousness during unresponsiveness induced by propofol, xenon, and ketamine. NeuroImage, 189, 631–644. https://doi.org/10.1016/j.neuroimage.2019.01.024

Comsa, I. M. (2019). Tracking brain dynamics across transitions of consciousness [Doctoral thesis]. University of Cambridge, Cambridge, England. https://doi.org/10.17863/CAM.37723

Curcio, G., Ferrara, M., Piergianni, A., Fratello, F., & Gennaro, L. de (2004). Paradoxes of the first-night effect: A quantitative analysis of antero-posterior EEG topography. Clinical Neurophysiology, 115(5), 1178–1188. https://doi.org/10.1016/j.clinph.2003.12.018

Dave, S., Brothers, T. A., & Swaab, T. Y. (2018). 1/f neural noise and electrophysiological indices of contextual prediction in aging. Brain Research, 1691, 34–43. https://doi.org/10.1016/j.brainres.2018.04.007

Davis, H., Davis, P. A., Loomis, A. L., Harvey, E. N., & Hobart, G. (1938). Human brain potentials during the onset of sleep. Journal of Neurophysiology, 1(1), 24–38.

Dijk, D.l.zJ. (1995). EEG slow waves and sleep spindles: windows on the sleeping brain. Behavioural Brain Research, 69(1-2), 109–116. https://doi.org/10.1016/0166-4328(95)00007-G

Donders, F. C. (1969). On the speed of mental processes. Acta Psychologica, 30, 412–431. https://doi.org/10.1016/0001-6918(69)90065-1

Donoghue, T., Haller, M., Peterson, E. J., Varma, P., Sebastian, P., Gao, R., Noto, T., Lara, A. H., Wallis, J. D., Knight, R. T., Shestyuk, A., & Voytek, B. (2020). Parameterizing neural power spectra into periodic and aperiodic components. Nature Neuroscience, 23(12), 1655–1665. https://doi.org/10.1038/s41593-020-00744-x

Ferenets, R., Vanluchene, A., Lipping, T., Heyse, B., & Struys, M. M. R. F. (2007). Behavior of entropy/complexity measures of the electroencephalogram during propofol-induced sedation: Dose-dependent effects of remifentanil. Anesthesiology, 106(4), 696–706. https://doi.org/10.1097/01.anes.0000264790.07231.2d

Figueiro, M. G., Sahin, L., Wood, B., & Plitnick, B. (2016). Light at Night and Measures of Alertness and Performance: Implications for Shift Workers. Biological Research for Nursing, 18(1), 90–100. https://doi.org/10.1177/1099800415572873

Friedrich, S., Konietschke, F., & Pauly, M. (2019). Resampling-Based Analysis of Multivariate Data and Repeated Measures Designs with the R Package MANOVA.RM. The R Journal, 11(2), 380–400. https://doi.org/10.32614/RJ-2019-051

Gao, R., & Penzes, P. (2015). Common mechanisms of excitatory and inhibitory imbalance in schizophrenia and autism spectrum disorders. Current Molecular Medicine, 15(2), 146–167. https://doi.org/10.2174/1566524015666150303003028

Gao, R., Peterson, E. J., & Voytek, B. (2017). Inferring synaptic excitation/inhibition balance from field potentials. NeuroImage, 158, 70–78. https://doi.org/10.1016/j.neuroimage.2017.06.078

Gerster, M., Waterstraat, G., Litvak, V., Lehnertz, K., Schnitzler, A., Florin, E., Curio, G., & Nikulin, V. (2022). Separating Neural Oscillations from Aperiodic 1/f Activity: Challenges and Recommendations. Neuroinformatics. Advance online publication. https://doi.org/10.1007/s12021-022-09581-8

González, J., Mateos, D., Cavelli, M., Mondino, A., Pascovich, C., Torterolo, P., & Rubido, N. (2022). Low frequency oscillations drive EEG’s complexity changes during wakefulness and sleep. Neuroscience, 494, 1–11. https://doi.org/10.1016/j.neuroscience.2022.04.025

Gratton, G., Coles, M. G., & Donchin, E. (1983). A new method for off-line removal of ocular artifact. Electroencephalography and Clinical Neurophysiology, 55(4), 468–484. https://doi.org/10.1016/0013-4694(83)90135-9

Halsband, U. (1998). Encoding and retrieval in declarative learning: a positron emission tomography study. Behavioural Brain Research, 97(1-2), 69–78. https://doi.org/10.1016/S0166-4328(98)00028-X

Harris, K. D., & Thiele, A. (2011). Cortical state and attention. Nature Reviews. Neuroscience, 12(9), 509–523. https://doi.org/10.1038/nrn3084

He, B. J. (2011). Scale-free properties of the functional magnetic resonance imaging signal during rest and task. The Journal of Neuroscience, 31(39), 13786–13795. https://doi.org/10.1523/JNEUROSCI.2111-11.2011

He, B. J. (2014). Scale-free brain activity: Past, present, and future. Trends in Cognitive Sciences, 18(9), 480–487. https://doi.org/10.1016/j.tics.2014.04.003

Höhn, C., Schmid, S. R., Plamberger, C. P., Bothe, K., Angerer, M., Gruber, G., Pletzer, B. A., & Hoedlmoser, K. (2021). Preliminary Results: The Impact of Smartphone Use and Short-Wavelength Light during the Evening on Circadian Rhythm, Sleep and Alertness. Clocks & Sleep, 3(1), 66–86. https://doi.org/10.3390/clockssleep3010005

Jacob, M. S., Roach, B. J., Sargent, K. S., Mathalon, D. H., & Ford, J. M. (2021). Aperiodic measures of neural excitability are associated with anticorrelated hemodynamic networks at rest: A combined EEG-fMRI study. NeuroImage, 245, 118705. https://doi.org/10.1016/j.neuroimage.2021.118705

Kanashiro, T., Ocker, G. K., Cohen, M. R., & Doiron, B. (2017). Attentional modulation of neuronal variability in circuit models of cortex. ELife, 6. https://doi.org/10.7554/eLife.23978

Karalunas, S. L., Ostlund, B. D., Alperin, B. R., Figuracion, M., Gustafsson, H. C., Deming, E. M., Foti, D., Antovich, D., Dude, J., Nigg, J., & Sullivan, E. (2022). Electroencephalogram aperiodic power spectral slope can be reliably measured and predicts ADHD risk in early development. Developmental Psychobiology, 64(3), e22228. https://doi.org/10.1002/dev.22228

Kirstein, C. (2007). Sleeping and Dreaming. In xPharm: The Comprehensive Pharmacology Reference (pp. 1–4). Elsevier. https://doi.org/10.1016/B978-008055232-3.60319-8

Kleiner, M., Brainard, D., Pelli, D., Ingling, A., & Murray, R., Broussard, C. (2007). What’s new in psychtoolbox-3. Perception, 36(14), 1–16.

Klimesch, W. (1999). EEG alpha and theta oscillations reflect cognitive and memory performance: a review and analysis. Brain Research Reviews, 29(2-3), 169–195. https://doi.org/10.1016/s0165-0173(98)00056-3

Klimesch, W., Schimke, H., & Pfurtscheller, G. (1993). Alpha frequency, cognitive load and memory performance. Brain Topography, 5(3), 241–251. https://doi.org/10.1007/BF01128991

Kozhemiako, N., Mylonas, D., Pan, J. Q., Prerau, M. J., Redline, S., & Purcell, S. M. (2022). Sources of variation in the spectral slope of the sleep EEG. ENeuro, 9(3). https://doi.org/10.1523/ENEURO.0094-22.2022

Kuula, L., & Pesonen, A.l.zK. (2021). Heart Rate Variability and Firstbeat Method for Detecting Sleep Stages in Healthy Young Adults: Feasibility Study. JMIR MHealth and UHealth, 9(2), e24704. https://doi.org/10.2196/24704

La Torre-Luque, A. de, Bornas, X., Balle, M., & Fiol-Veny, A. (2016). Complexity and nonlinear biomarkers in emotional disorders: A meta-analytic study. Neuroscience and Biobehavioral Reviews, 68, 410–422. https://doi.org/10.1016/j.neubiorev.2016.05.023

Lau, Z. J., Pham, T., Chen, S. H. A., & Makowski, D. (2022). Brain entropy, fractal dimensions and predictability: A review of complexity measures for EEG in healthy and neuropsychiatric populations. The European Journal of Neuroscience. Advance online publication. https://doi.org/10.1111/ejn.15800

Lempel, A., & Ziv, J. (1976). On the Complexity of Finite Sequences. IEEE Transactions on Information Theory, 22(1), 75–81. https://doi.org/10.1109/TIT.1976.1055501

Lendner, J. D., Helfrich, R. F., Mander, B. A., Romundstad, L., Lin, J. J., Walker, M. P., Larsson, P. G., & Knight, R. T. (2020). An electrophysiological marker of arousal level in humans. ELife, 9. https://doi.org/10.7554/eLife.55092

Lendner, J. D., Mander, B. A., Schuh-Hofer, S., Schmidt, H., Knight, R. T., Walker, M. P., Lin, J. J., & Helfrich, R. F. (2022). Human REM sleep controls neural excitability in support of memory formation. BioRxiv. Advance online publication. https://doi.org/10.1101/2022.05.13.491801

Lüdtke, S., Hermann, W., Kirste, T., Beneš, H., & Teipel, S. (2021). An algorithm for actigraphy-based sleep/wake scoring: Comparison with polysomnography. Clinical Neurophysiology, 132(1), 137–145. https://doi.org/10.1016/j.clinph.2020.10.019

Ma, Y., Shi, W., Peng, C.l.zK., & Yang, A. C. (2018). Nonlinear dynamical analysis of sleep electroencephalography using fractal and entropy approaches. Sleep Medicine Reviews, 37, 85–93. https://doi.org/10.1016/j.smrv.2017.01.003

Mateos, D., Guevara Erra, R., Wennberg, R., & Perez Velazquez, J. L. (2018). Measures of entropy and complexity in altered states of consciousness. Cognitive Neurodynamics, 12(1), 73–84. https://doi.org/10.1007/s11571-017-9459-8

Medel, V., Irani, M., Ossandón, T., & Boncompte, G. (2020). Complexity and 1/f slope jointly reflect cortical states across different E/I balances. BioRxiv. Advance online publication. https://doi.org/10.1101/2020.09.15.298497

Mediano, P. A. M., Ikkala, A., Kievit, R. A., Jagannathan, S. R., Varley, T. F., Stamatakis, E. A., Bekinschtein, T. A., & Bor, D. (2021). Fluctuations in Neural Complexity During Wakefulness Relate To Conscious Level and Cognition. BioRxiv. Advance online publication. https://doi.org/10.1101/2021.09.23.461002

Miskovic, V., MacDonald, K. J., Rhodes, L. J., & Cote, K. A. (2019). Changes in EEG multiscale entropy and power-law frequency scaling during the human sleep cycle. Human Brain Mapping, 40(2), 538–551. https://doi.org/10.1002/hbm.24393

Mitsukura, Y., Fukunaga, K., Yasui, M., & Mimura, M. (2020). Sleep stage detection using only heart rate. Health Informatics Journal, 26(1), 376–387. https://doi.org/10.1177/1460458219827349

Oostenveld, R., Fries, P., Maris, E., & Schoffelen, J.l.zM. (2011). Fieldtrip: Open source software for advanced analysis of MEG, EEG, and invasive electrophysiological data. Computational Intelligence and Neuroscience, 2011, 156869. https://doi.org/10.1155/2011/156869

Ouyang, G., Hildebrandt, A., Schmitz, F., & Herrmann, C. S. (2020). Decomposing alpha and 1/f brain activities reveals their differential associations with cognitive processing speed. NeuroImage, 205, 116304. https://doi.org/10.1016/j.neuroimage.2019.116304

Pascovich, C., Castro-Zaballa, S., Mediano, P. A. M., Bor, D., Canales-Johnson, A., Torterolo, P., & Bekinschtein, T. A. (2022). Ketamine and sleep modulate neural complexity dynamics in cats. The European Journal of Neuroscience, 55(6), 1584–1600. https://doi.org/10.1111/ejn.15646

Pathania, A., Euler, M. J., Clark, M., Cowan, R. L., Duff, K., & Lohse, K. R. (2022). Resting EEG spectral slopes are associated with age-related differences in information processing speed. Biological Psychology, 168, 108261. https://doi.org/10.1016/j.biopsycho.2022.108261

Peever, J., & Fuller, P. M. (2017). The Biology of REM Sleep. Current Biology, 27(22), R1237–R1248. https://doi.org/10.1016/j.cub.2017.10.026

Peigneux, P., Laureys, S., Fuchs, S., Delbeuck, X., Degueldre, C., Aerts, J., Delfiore, G., Luxen, A., & Maquet, P. (2001). Generation of rapid eye movements during paradoxical sleep in humans. NeuroImage, 14(3), 701–708. https://doi.org/10.1006/nimg.2001.0874

Pereda, E., Gamundi, A., Rial, R., & González, J. (1998). Non-linear behaviour of human EEG: fractal exponent versus correlation dimension in awake and sleep stages. Neuroscience Letters, 250(2), 91–94. https://doi.org/10.1016/S0304-3940(98)00435-2

Plamberger, C. P., van Wijk, H. E., Kerschbaum, H., Pletzer, B. A., Gruber, G., Oberascher, K., Dresler, M., Hahn, M. A., & Hoedlmoser, K. (2021). Impact of menstrual cycle phase and oral contraceptives on sleep and overnight memory consolidation. Journal of Sleep Research, 30(4), e13239. https://doi.org/10.1111/jsr.13239

Podvalny, E., Noy, N., Harel, M., Bickel, S., Chechik, G., Schroeder, C. E., Mehta, A. D., Tsodyks, M., & Malach, R. (2015). A unifying principle underlying the extracellular field potential spectral responses in the human cortex. Journal of Neurophysiology, 114(1), 505–519. https://doi.org/10.1152/jn.00943.2014

Radha, M., Fonseca, P., Moreau, A., Ross, M., Cerny, A., Anderer, P., Long, X., & Aarts, R. M. (2019). Sleep stage classification from heart-rate variability using long short-term memory neural networks. Scientific Reports, 9(1), 14149. https://doi.org/10.1038/s41598-019-49703-y

Richard, B. B., Albertario, C. L., Harding, S. M., Uoyd, R. M., Plante, D. T., Quan, S. F., Troester, M. M., & Vaughn, B. V. (2012). The AASM Manual for the Scoring of Sleep and Associated Events: Rules, Terminology and Technical Specifications (2nd ed.). American Academy of Sleep Medicine: Darien, IL. www.aasmnet.org

Rivolta, M. W., Migliorini, M., Aktaruzzaman, M., Sassi, R., & Bianchi, A. M. (2014). Effects of the series length on Lempel-Ziv Complexity during sleep. Annual International Conference of the IEEE Engineering in Medicine and Biology Society. IEEE Engineering in Medicine and Biology Society. Annual International Conference, 2014, 693–696. https://doi.org/10.1109/EMBC.2014.6943685

Roberts, D. M., Schade, M. M., Mathew, G. M., Gartenberg, D., & Buxton, O. M. (2020). Detecting sleep using heart rate and motion data from multisensor consumer-grade wearables, relative to wrist actigraphy and polysomnography. Sleep, 43(7). https://doi.org/10.1093/sleep/zsaa045

Robertson, M. M., Furlong, S., Voytek, B., Donoghue, T., Boettiger, C. A., & Sheridan, M. A. (2019). Eeg power spectral slope differs by ADHD status and stimulant medication exposure in early childhood. Journal of Neurophysiology, 122(6), 2427–2437. https://doi.org/10.1152/jn.00388.2019

Rubenstein, J. L. R., & Merzenich, M. M. (2003). Model of autism: Increased ratio of excitation/inhibition in key neural systems. Genes, Brain, and Behavior, 2(5), 255–267. https://doi.org/10.1034/j.1601-183X.2003.00037.x

Sadeh, A., Alster, J., Urbach, D., & Lavie, P. (1989). Actigraphically based automatic bedtime sleep-wake scoring: Validity and clinical applications. Journal of Ambulatory Monitoring(3), 209–216.

Schartner, M., Pigorini, A., Gibbs, S. A., Arnulfo, G., Sarasso, S., Barnett, L., Nobili, L., Massimini, M., Seth, A., & Barrett, A. B. (2017). Global and local complexity of intracranial EEG decreases during NREM sleep. Neuroscience of Consciousness, 2017(1), 1-12. https://doi.org/10.1093/nc/niw022

Schartner, M., Seth, A., Noirhomme, Q., Boly, M., Bruno, M.l.zA., Laureys, S., & Barrett, A. B. (2015). Complexity of Multi-Dimensional Spontaneous EEG Decreases during Propofol Induced General Anaesthesia. PloS One, 10(8), e0133532. https://doi.org/10.1371/journal.pone.0133532

Schaworonkow, N., & Voytek, B. (2021). Longitudinal changes in aperiodic and periodic activity in electrophysiological recordings in the first seven months of life. Developmental Cognitive Neuroscience, 47, 100895. https://doi.org/10.1016/j.dcn.2020.100895

Schmid, S. R., Höhn, C., Bothe, K., Plamberger, C. P., Angerer, M., Pletzer, B. A., & Hoedlmoser, K. (2021). How Smart Is It to Go to Bed with the Phone? The Impact of Short-Wavelength Light and Affective States on Sleep and Circadian Rhythms. Clocks & Sleep, 3(4), 558–580. https://doi.org/10.3390/clockssleep3040040

Sheehan, T. C., Sreekumar, V., Inati, S. K., & Zaghloul, K. A. (2018). Signal Complexity of Human Intracranial EEG Tracks Successful Associative-Memory Formation across Individuals. The Journal of Neuroscience, 38(7), 1744–1755. https://doi.org/10.1523/JNEUROSCI.2389-17.2017

Siegel, J. M. (2011). Rem sleep: A biological and psychological paradox. Sleep Medicine Reviews, 15(3), 139–142. https://doi.org/10.1016/j.smrv.2011.01.001

Sweller, J. (2011). Cognitive Load Theory. In Psychology of Learning and Motivation (Vol. 55, pp. 37– 76). Elsevier. https://doi.org/10.1016/B978-0-12-387691-1.00002-8

Symonds, C. (1959). Excitation and inhibition in epilepsy. Proceedings of the Royal Society of Medicine, 52(6), 395–402.

Tal, A., Shinar, Z., Shaki, D., Codish, S., & Goldbart, A. (2017). Validation of Contact-Free Sleep Monitoring Device with Comparison to Polysomnography. Journal of Clinical Sleep Medicine, 13(3), 517–522. https://doi.org/10.5664/jcsm.6514

Terzano, M. G., Parrino, L., Smerieri, A., Chervin, R., Chokroverty, S., Guilleminault, C., Hirshkowitz, M., Mahowald, M., Moldofsky, H., Rosa, A., Thomas, R., & Walters, A. (2002). Atlas, rules, and recording techniques for the scoring of cyclic alternating pattern (CAP) in human sleep. Sleep Medicine, 3(2), 187–199. https://doi.org/10.1016/S1389-9457(01)00149-6

Tosun, P. D., Dijk, D.l.zJ., Winsky-Sommerer, R., & Abásolo, D. (2019). Effects of Ageing and Sex on Complexity in the Human Sleep EEG: A Comparison of Three Symbolic Dynamic Analysis Methods. Complexity, 2019, 1–12. https://doi.org/10.1155/2019/9254309

Treder, M. S. (2020). Mvpa-Light: A Classification and Regression Toolbox for Multi-Dimensional Data. Frontiers in Neuroscience, 14, 289. https://doi.org/10.3389/fnins.2020.00289

van de Borne, P., Nguyen, H., Biston, P., Linkowski, P., & Degaute, J. P. (1994). Effects of wake and sleep stages on the 24-h autonomic control of blood pressure and heart rate in recumbent men. The American Journal of Physiology, 266(2), 548–554. https://doi.org/10.1152/ajpheart.1994.2662.H548

Voytek, B., & Knight, R. T. (2015). Dynamic network communication as a unifying neural basis for cognition, development, aging, and disease. Biological Psychiatry, 77(12), 1089–1097. https://doi.org/10.1016/j.biopsych.2015.04.016

Voytek, B., Kramer, M. A., Case, J., Lepage, K. Q., Tempesta, Z. R., Knight, R. T., & Gazzaley, A. (2015). Age-Related Changes in 1/f Neural Electrophysiological Noise. The Journal of Neuroscience, 35(38), 13257–13265. https://doi.org/10.1523/JNEUROSCI.2332-14.2015

Waschke, L., Donoghue, T., Fiedler, L., Smith, S., Garrett, D. D., Voytek, B., & Obleser, J. (2021). Modality-specific tracking of attention and sensory statistics in the human electrophysiological spectral exponent. ELife, 10. https://doi.org/10.7554/eLife.70068

Waschke, L., Wöstmann, M., & Obleser, J. (2017). States and traits of neural irregularity in the age-varying human brain. Scientific Reports, 7(1), 17381. https://doi.org/10.1038/s41598-017-17766-4

Welch (1984). A Technique for High-Performance Data Compression. Computer, 17(6), 8–19. https://doi.org/10.1109/MC.1984.1659158

Wickham, H. (2016). Ggplot2: Elegant graphics for data analysis (Second edition). Use R! Springer. https://doi.org/10.1007/978-3-319-24277-4

Wong, M. (2010). Too much inhibition leads to excitation in absence epilepsy. Epilepsy Currents, 10(5), 131–132. https://doi.org/10.1111/j.1535-7511.2010.01379.x

Zhang, X. S., Roy, R. J., & Jensen, E. W. (2001). Eeg complexity as a measure of depth of anesthesia for patients. IEEE Transactions on Bio-Medical Engineering, 48(12), 1424–1433. https://doi.org/10.1109/10.966601

Zhu, X., Xu, H., Zhao, J., & Tian, J. (2017). Automated Epileptic Seizure Detection in Scalp EEG Based on Spatial-Temporal Complexity. Complexity, 2017, 1–8. https://doi.org/10.1155/2017/5674392

