## Supplementary File for "Spectral slope and Lempel-Ziv complexity as robust markers of brain states during sleep and wakefulness"

**Table 1 related to Figure 1.** Questionnaires from the entrance examination and sum-scores (mean ± standard deviation) for the analyzed sample as well as recommended cut-off values for clinical or extreme populations (*N* = 28).

| Questionnaire | Mean (SD) | Cut-Off |
| --- | --- | --- |
| Pittsburgh Sleep Quality Index (Buysse et al., 1989) | 3.36 (2.11) | ≥ 10 |
| State Trait Anxiety Inventory: Trait (Spielberger et al., 1970) | 33.39 (8.39) | ≥ 45 |
| Social Interaction Anxiety Scale (Mattick & Clarke, 1998) | 23.28 (15.75) | ≥ 30 |
| Beck-Depression-Inventory (Hautzinger et al., 2009) | 3.96 (4.44) | ≥ 18 |
| Perceived Stress Scale (Cohen et al., 1983) | 10.36 (5.14) | ≥ 27 |
| Smartphone Addiction Scale (Kwon et al., 2013) | 20.07 (5.70) | ≥ 31 |
| Morning-Eveningness Questionnaire (Griefahn et al., 2001) | 53.96 (8.58) | ≤ 30 or ≥ 70 |

**Note.** Cut-Off values were taken from the information provided in the respective manual. Recommendations from Stojanović et al. (2020) were used to obtain a cut-off value for the trait version of the State Trait Anxiety Inventory (STAI-T).

**Table 2: Classifier performance across all sleep stages and tasks, 30 – 45Hz.** Classification accuracy for all pairwise combinations of sleep stage and task. *Upper triangular matrix shows the results for Lempel-Ziv complexity* and l*ower triangular matrix for the spectral slope*. The data was pooled over all lab-visits for each subject (*N* = 28).

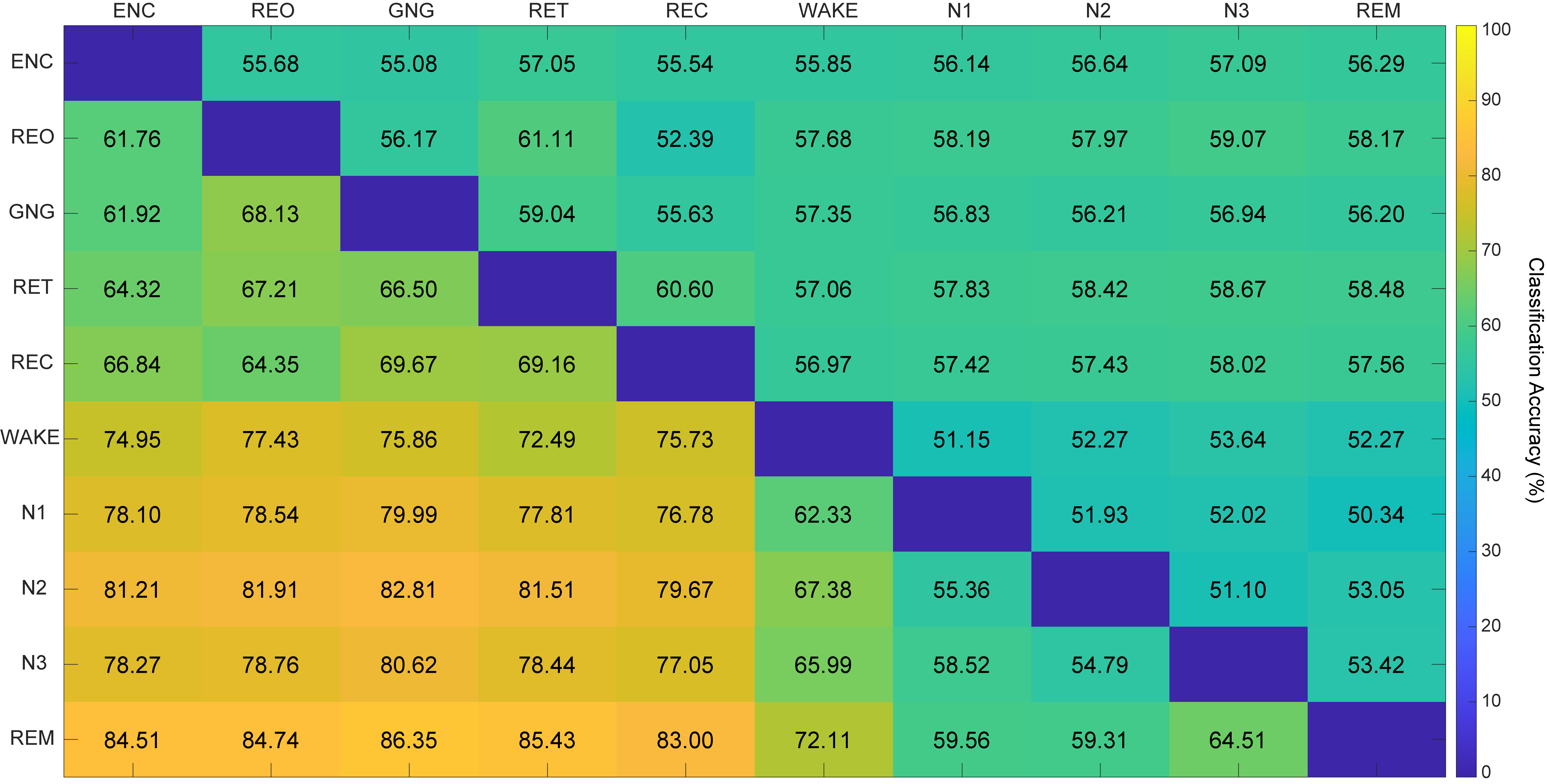

**Table 3: Classifier performance across all sleep stages and tasks, 1 – 45Hz.** Classification accuracy for all pairwise combinations of sleep stage and task. *Upper triangular matrix shows the results for Lempel-Ziv complexity* and the l*ower triangular matrix for the spectral slope*. The data was pooled over all lab-visits for each subject (*N* = 28).

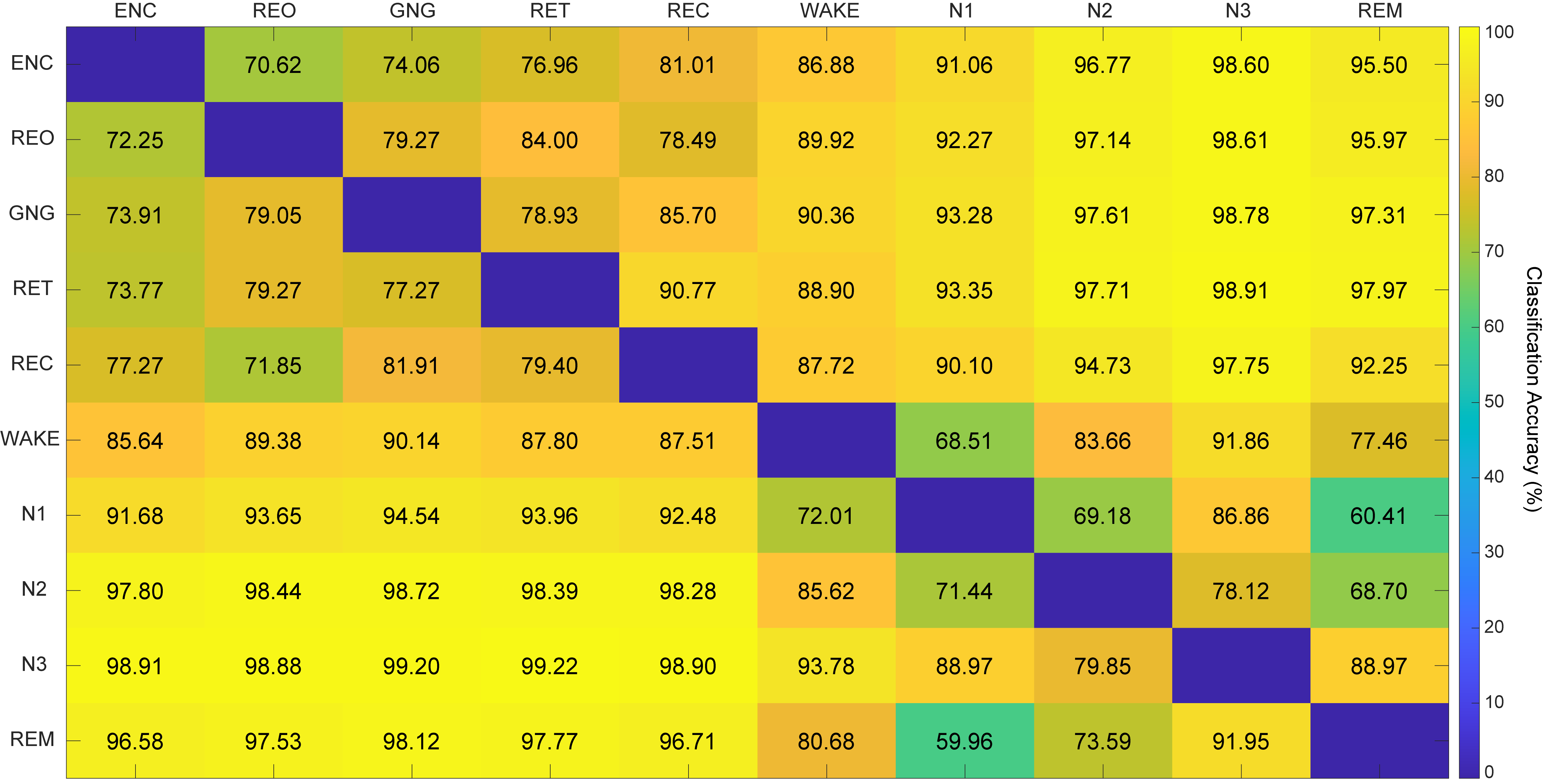

**Table 4: Robustness of the spectral slope and Lempel-Ziv complexity across lab-visits.** Correlation coefficients over all electrodes for each parameter between the three experimental recordings (1 x 2, 1 x 3 and 2 x 3). Each of the experimental recordings involved a different light condition (cf., Höhn et al., 2021) and refers to one of the three lab-visits per subject. Spearman rho correlation coefficients are shown when the assumption of normality was violated.

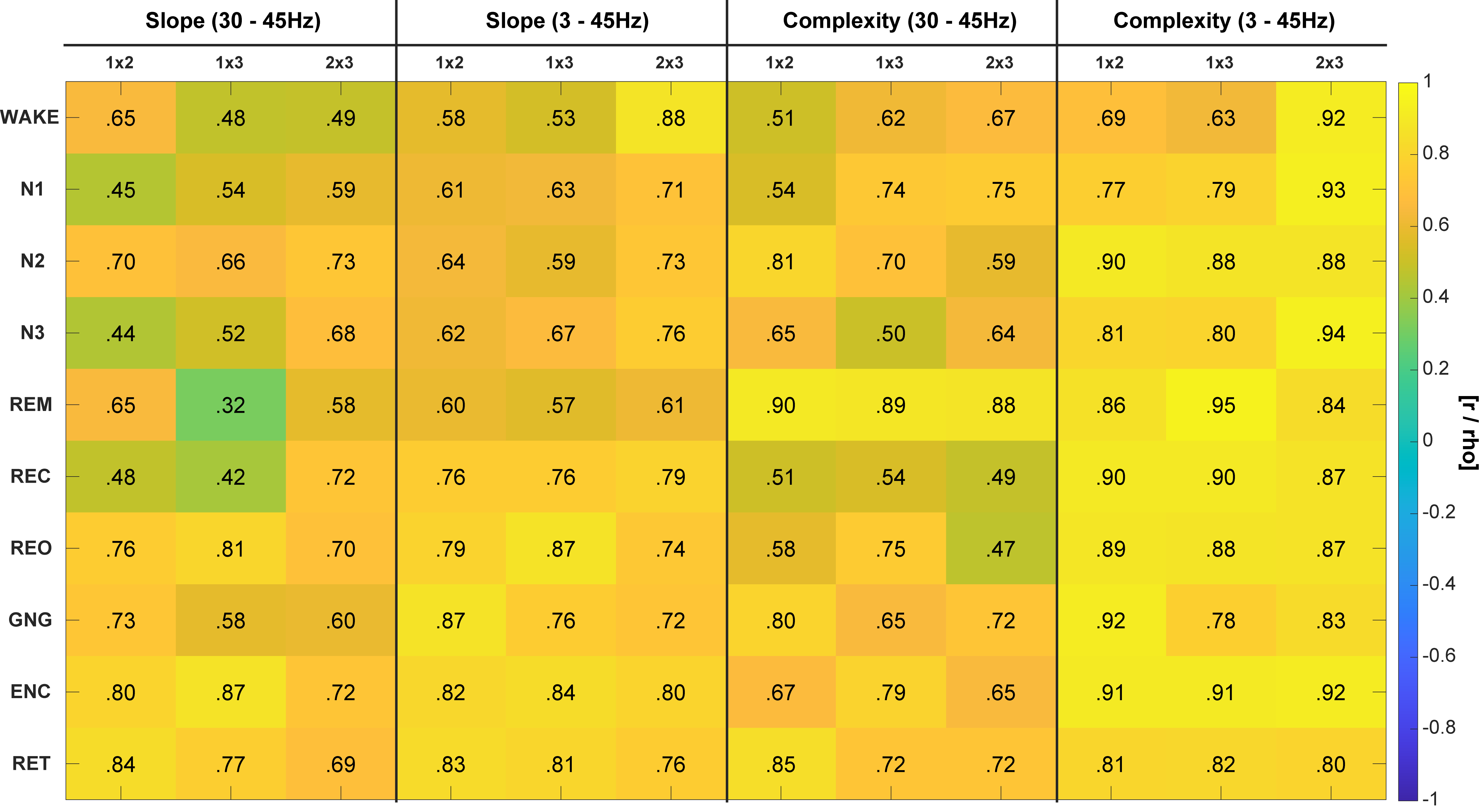

**Table 5: Descriptive overview of the sleep architecture for each night.** Whole night sleep architecture for all lab visits (median and interquartile range; *N* = 28). TIB = Time in bed, TST = Total sleep time, SEFF = Sleep efficiency, SOL N2 = Sleep onset latency to N2, WASO = Wake time after sleep onset.

|  | Adapt | Exp. Recording #1 | Exp. Recording #2 | Exp. Recording #3 |
| --- | --- | --- | --- | --- |
| TIB (min) | 480.75 (1.00) | 480.50 (0.13) | 480.50 (0.50) | 480.50 (0.50) |
| TST (min) | 461 (23.88) | 466 (27.63) | 464.25 (18.75) | 468.50 (15.75) |
| SEFF (%) | 95.85 (4.99) | 97.14 (5.18) | 96.83 (4.08) | 97.50 (2.91) |
| SOL N2 (min) | 17.50 (12.25) | 12 (6.88) | 11.25 (9.38) | 11 (6.82) |
| N1 (%) | 15.45 (6.82) | 12.77 (7.00) | 10.40 (6.14) | 10.59 (6.14) |
| N2 (%) | 40.69 (12.27) | 38.32 (8.32) | 39.38 (8.07) | 39.01 (7.12) |
| N3 (%) | 27.43 (10.45) | 28.13 (11.64) | 29.28 (6.72) | 29.32 (8.69) |
| REM (%) | 15.46 (6.45) | 19.58 (7.27) | 20.43 (7.88) | 20.10 (5.58) |
| WASO (min) | 11.50 (25.75) | 9 (13.5) | 10.25 (11.88) | 7.75 (10.25) |

**Table 6: EEG preprocessing resulting number of epochs / segments.** Mean number of clean epochs (min, max) for all tasks and sleep stages per experimental condition (i.e., different lab-visits; for details see Höhn et al., 2021 and Schmid et al., 2021). For the wakefulness recordings, the data is averaged over the multiple measurements per lab-visit and the encoding session has been pooled over both runs per visit (*N* = 28).

| **A) Epochs for the multivariate pattern analyses (MVPA)** | | | |
| --- | --- | --- | --- |
| Task / Sleep-Stage | Exp. Recording #1 | Exp. Recording #2 | Exp. Recording #3 |
| Resting eyes closed | 42 (36, 45) | 41 (27, 45) | 42 (26, 45) |
| Resting eyes open | 41 (34, 45) | 39 (16, 45) | 41 (30, 45) |
| Go/Nogo task | 45 (40, 45) | 45 (39, 45) | 45 (45, 45) |
| Encoding | 46 (46, 46) | 46 (46, 46) | 46 (46, 46) |
| Retrieval | 43 (19, 45) | 43 (24, 45) | 43 (20, 45) |
| Wake (stage) | 45 (45, 45) | 45 (45, 45) | 45 (44, 45) |
| NREM1-3 & REM | 45 (45, 45) | 45 (45, 45) | 45 (45, 45) |
| **B) Epochs for all other analyses** | | | |
| Task / Sleep-Stage | Exp. Recording #1 | Exp. Recording #2 | Exp. Recording #3 |
| Resting eyes closed | 43 (36, 53) | 42 (27, 52) | 43 (26, 48) |
| Resting eyes open | 42 (34, 48) | 39 (16, 46) | 41 (30, 47) |
| Go/Nogo task | 143 (55, 165) | 136 (91, 165) | 138 (91, 161) |
| Encoding | 307 (65, 400) | 302 (168, 402) | 324 (208, 409) |
| Retrieval | 96 (19, 191) | 90 (24, 145) | 96 (20, 171) |
| Wake (stage) | 353 (47, 2099) | 301 (71, 1735) | 212 (44, 616) |
| NREM1 | 843 (395, 1748) | 848 (335, 2526) | 821 (415, 1453) |
| NREM2 | 2614 (1212, 3387) | 2585 (1244, 3339) | 2688 (1893, 3417) |
| NREM3 | 1916 (1109, 3220) | 1939 (1032, 3500) | 1966 (1233, 2940) |
| REM | 1224 (395, 1981) | 1312 (523, 1918) | 1327 (540, 1946) |

References

Buysse, D. J., Reynolds, C. F., Monk, T. H., Berman, S. R., & Kupfer, D. J. (1989). The Pittsburgh sleep quality index: A new instrument for psychiatric practice and research. *Psychiatry Research*, *28*(2), 193–213. https://doi.org/10.1016/0165-1781(89)90047-4

Cohen, S., Kamarck, T., & Mermelstein, R. (1983). A Global Measure of Perceived Stress. *Journal of Health and Social Behavior*, *24*(4), 385. https://doi.org/10.2307/2136404

Griefahn, B., Kunemund, C., Brode, P., & Mehnert, P. (2001). Zur Validität der deutschen Übersetzung des Morningness-Eveningness-Questionnaires von Horne und Ostberg. The Validity of a German Version of the Morningness-Eveningness-Questionnaire Developed by Horne and Ostberg. *Somnologie*, *5*(2), 71–80. https://doi.org/10.1046/j.1439-054X.2001.01149.x

Hautzinger, M., Keller, F., & Kühner, C. (2009). *Beck Depressions-Inventar (BDI-II)*. Revision. Frankfurt am Main: Pearson Assessment (2nd ed.).

Höhn, C., Schmid, S. R., Plamberger, C. P., Bothe, K., Angerer, M., Gruber, G., Pletzer, B. A., & Hoedlmoser, K. (2021). Preliminary Results: The Impact of Smartphone Use and Short-Wavelength Light during the Evening on Circadian Rhythm, Sleep and Alertness. *Clocks & Sleep*, *3*(1), 66–86. https://doi.org/10.3390/clockssleep3010005

Kwon, M., Kim, D.‑J., Cho, H., & Yang, S. (2013). The smartphone addiction scale: Development and validation of a short version for adolescents. *PloS One*, *8*(12), e83558. https://doi.org/10.1371/journal.pone.0083558

Mattick, R. P., & Clarke, J. C. (1998). Development and validation of measures of social phobia scrutiny fear and social interaction anxiety. *Behaviour Research and Therapy*, *36*(4), 455–470. https://doi.org/10.1016/S0005-7967(97)10031-6

Schmid, S. R., Höhn, C., Bothe, K., Plamberger, C. P., Angerer, M., Pletzer, B. A., & Hoedlmoser, K. (2021). How Smart Is It to Go to Bed with the Phone? The Impact of Short-Wavelength Light and Affective States on Sleep and Circadian Rhythms. *Clocks & Sleep*, *3*(4), 558–580. https://doi.org/10.3390/clockssleep3040040

Spielberger, C. D., Gorsuch, R. L., & Lushene, R. H. (1970). *Manual for the State-Trait Anxiety Inventory*. Palo Alto, CA: Consulting Psychologists Press, Inc.

Stojanović, N., Ranđelović, P., Nikolić, G., Stojiljković, N., Ilić, S., Stoiljković, B., & Radulović, N. (2020). Reliability and validity of the Spielberger's State-Trait Anxiety Inventory (STAI) in Serbian university student and psychiatric non-psychotic outpatient populations. *Acta Facultatis Medicae Naissensis*, *37*(2), 149–159. https://doi.org/10.5937/afmnai37-25011
