## Supplementary figures and images for "Spectral slope and Lempel-Ziv complexity as robust markers of brain states during sleep and wakefulness"

### Figure1_sup1.pdf

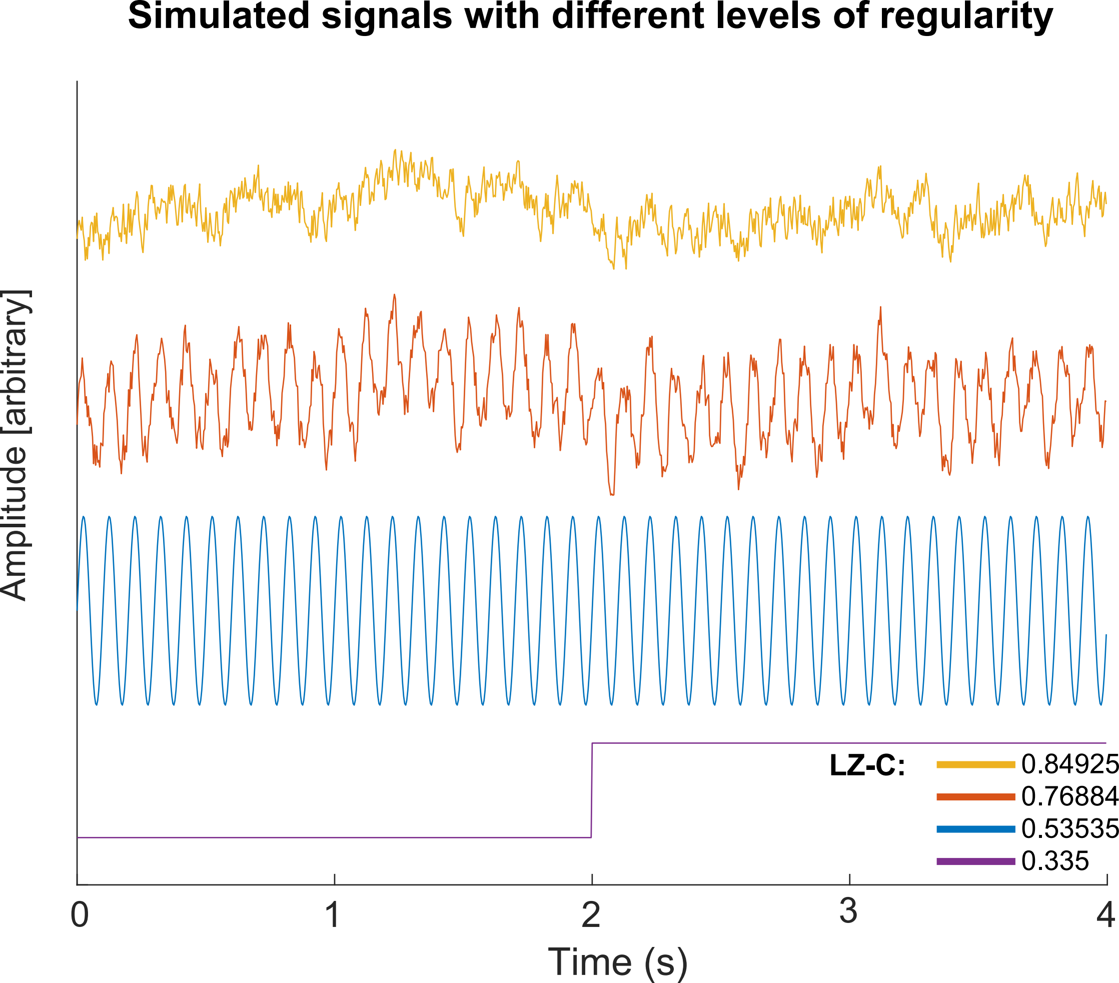

### Figure2_supp1.pdf

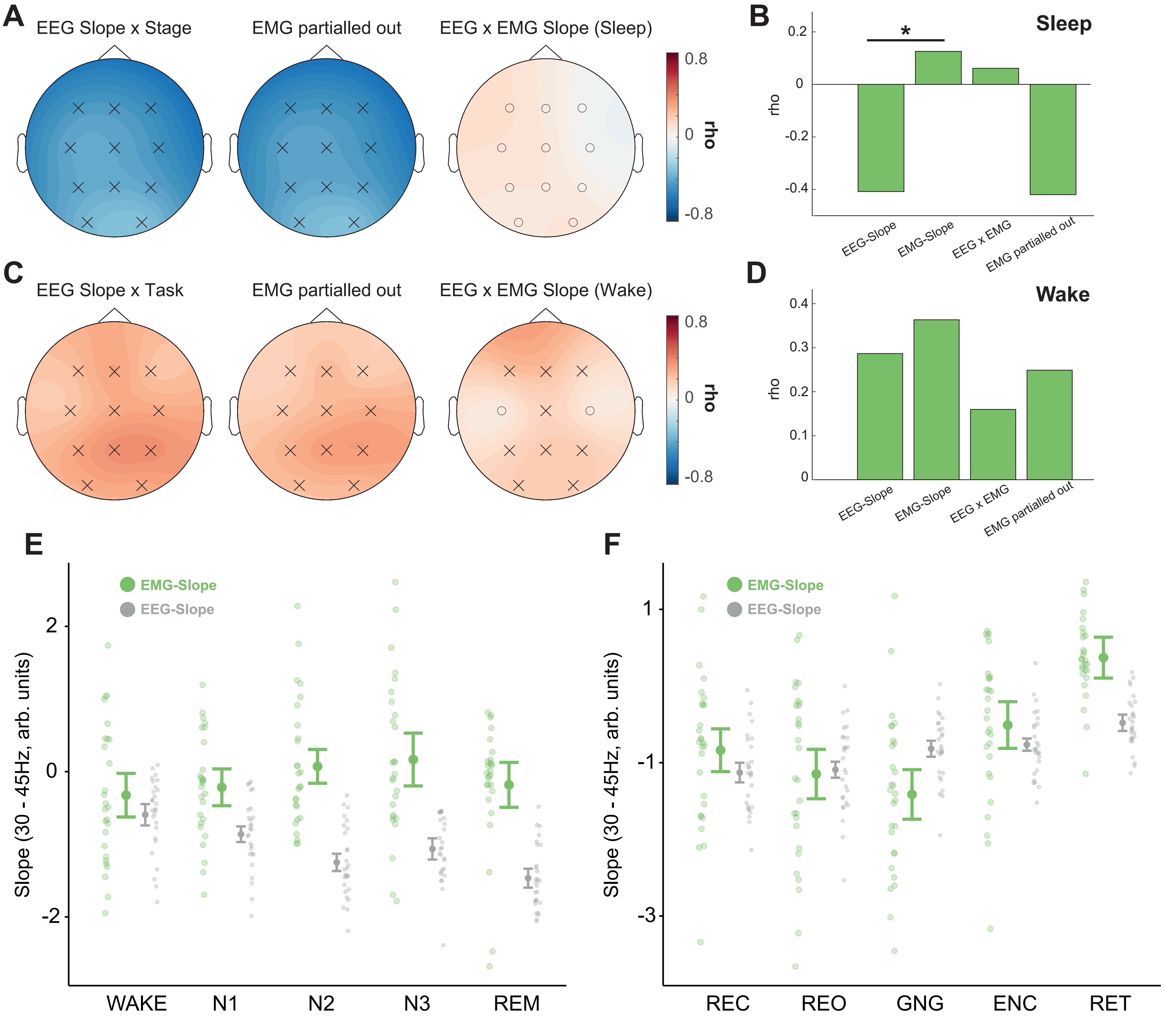

### Figure2_supp2.pdf

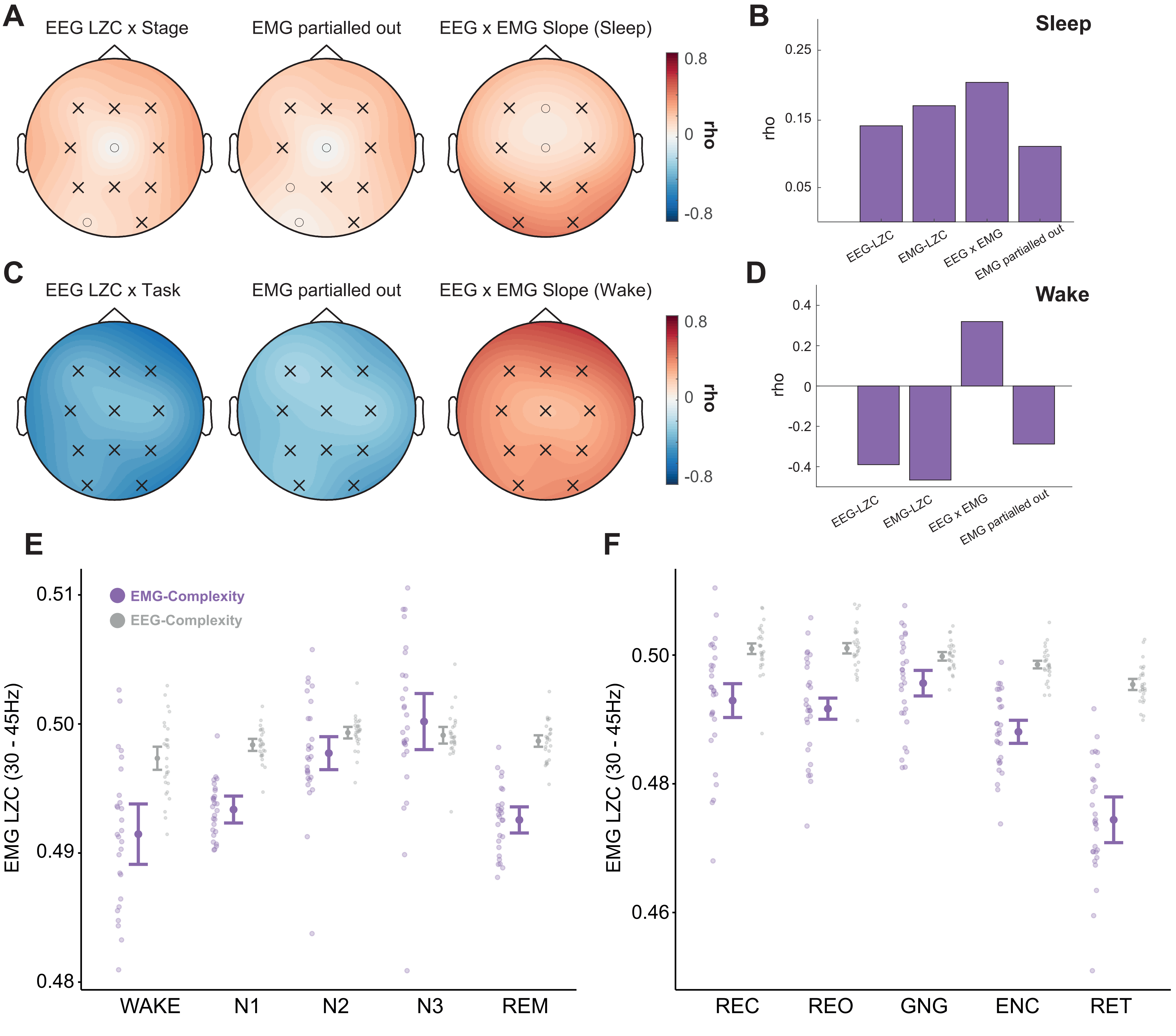

### Figure5_sup1.pdf

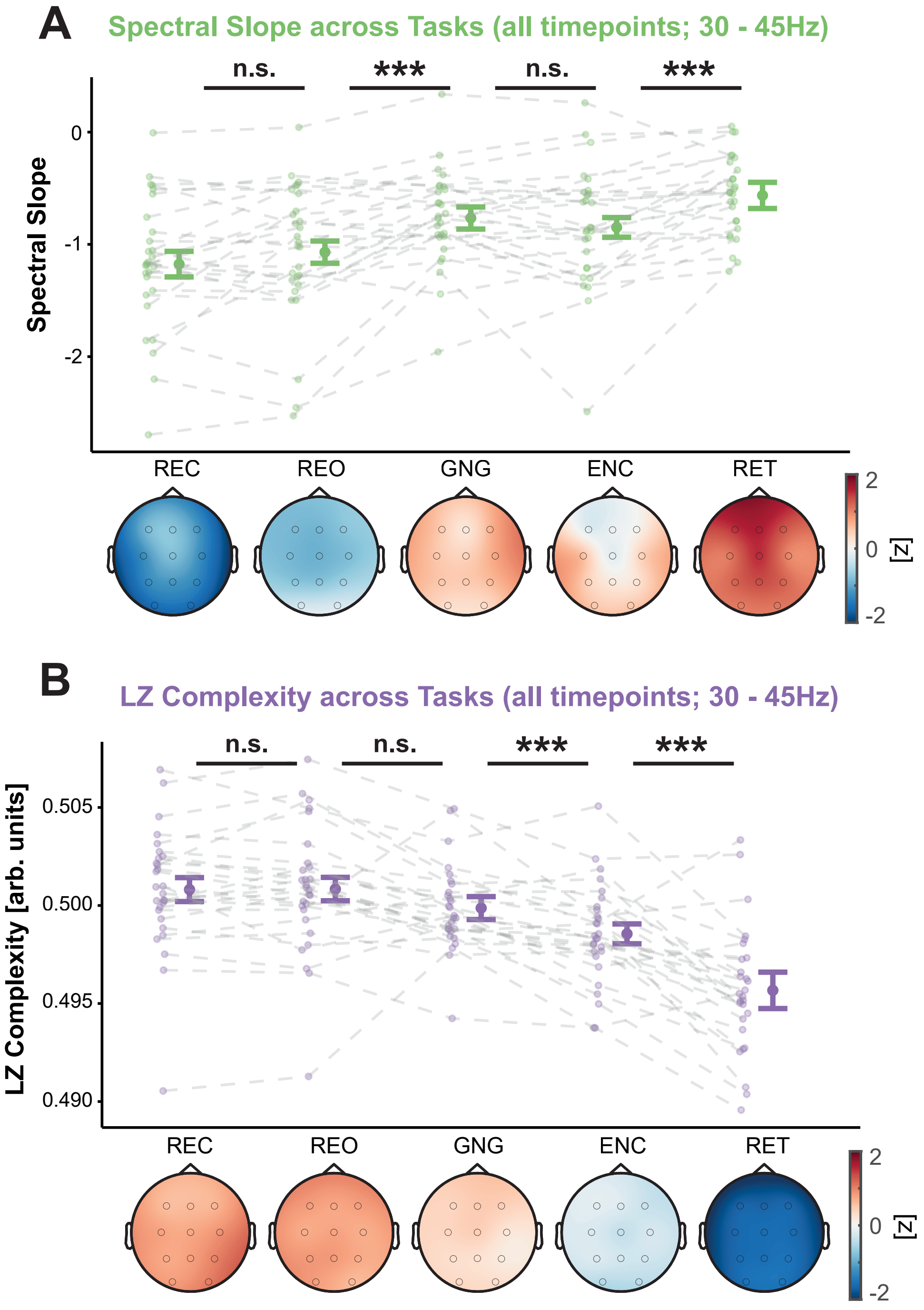

### Figure5_sup2.pdf

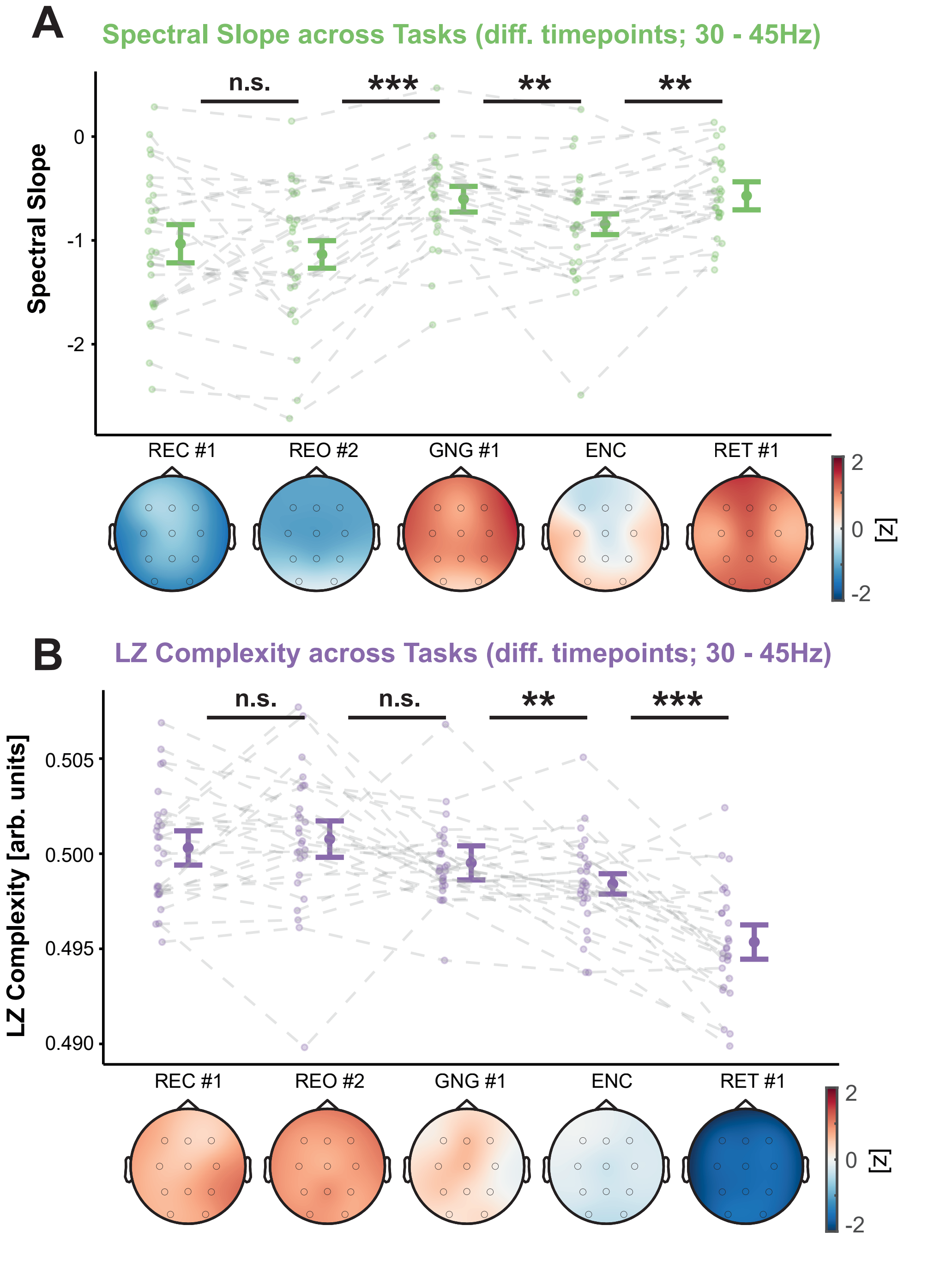

### Figure6_sup1.pdf

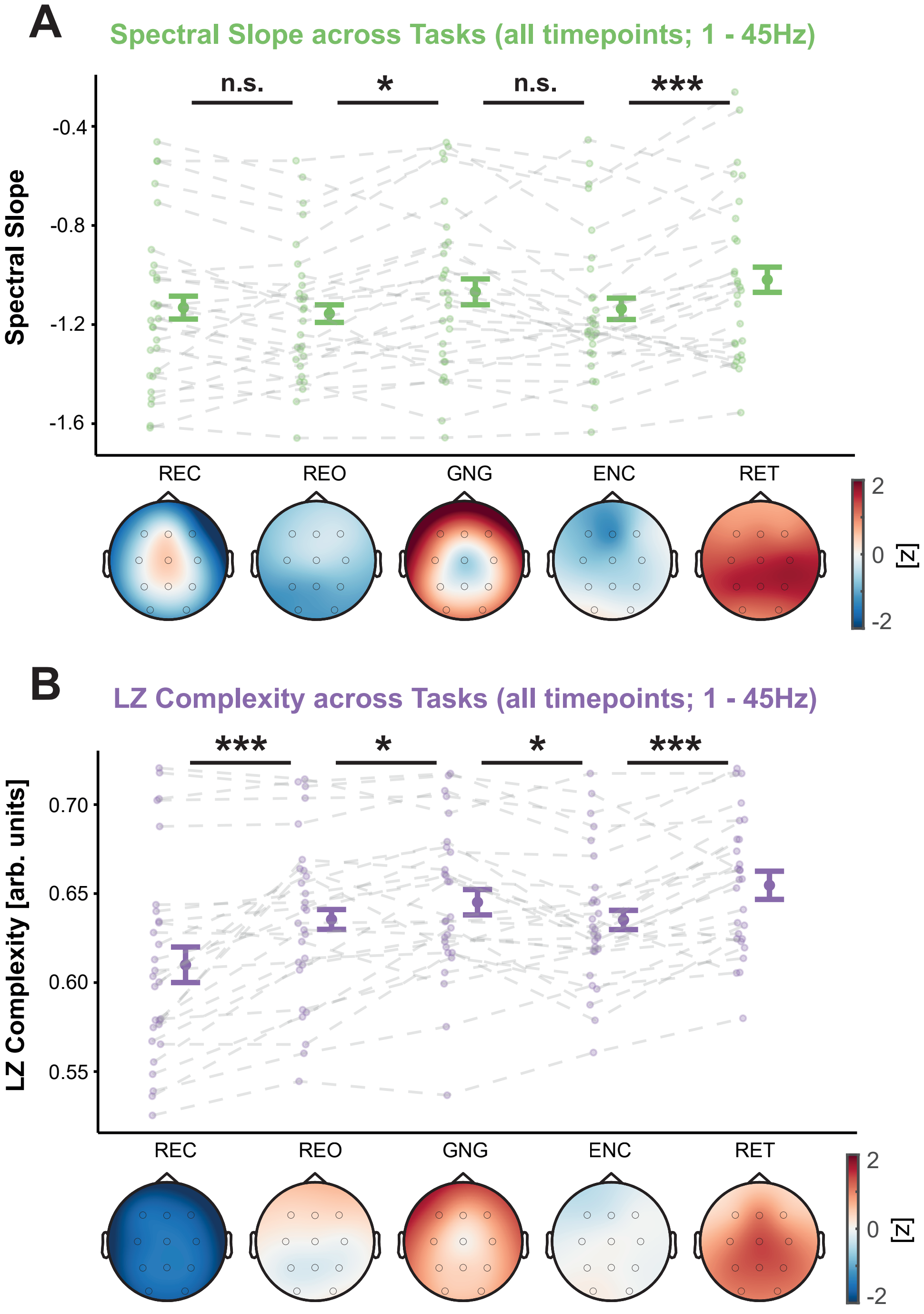

### Figure6_sup2.pdf

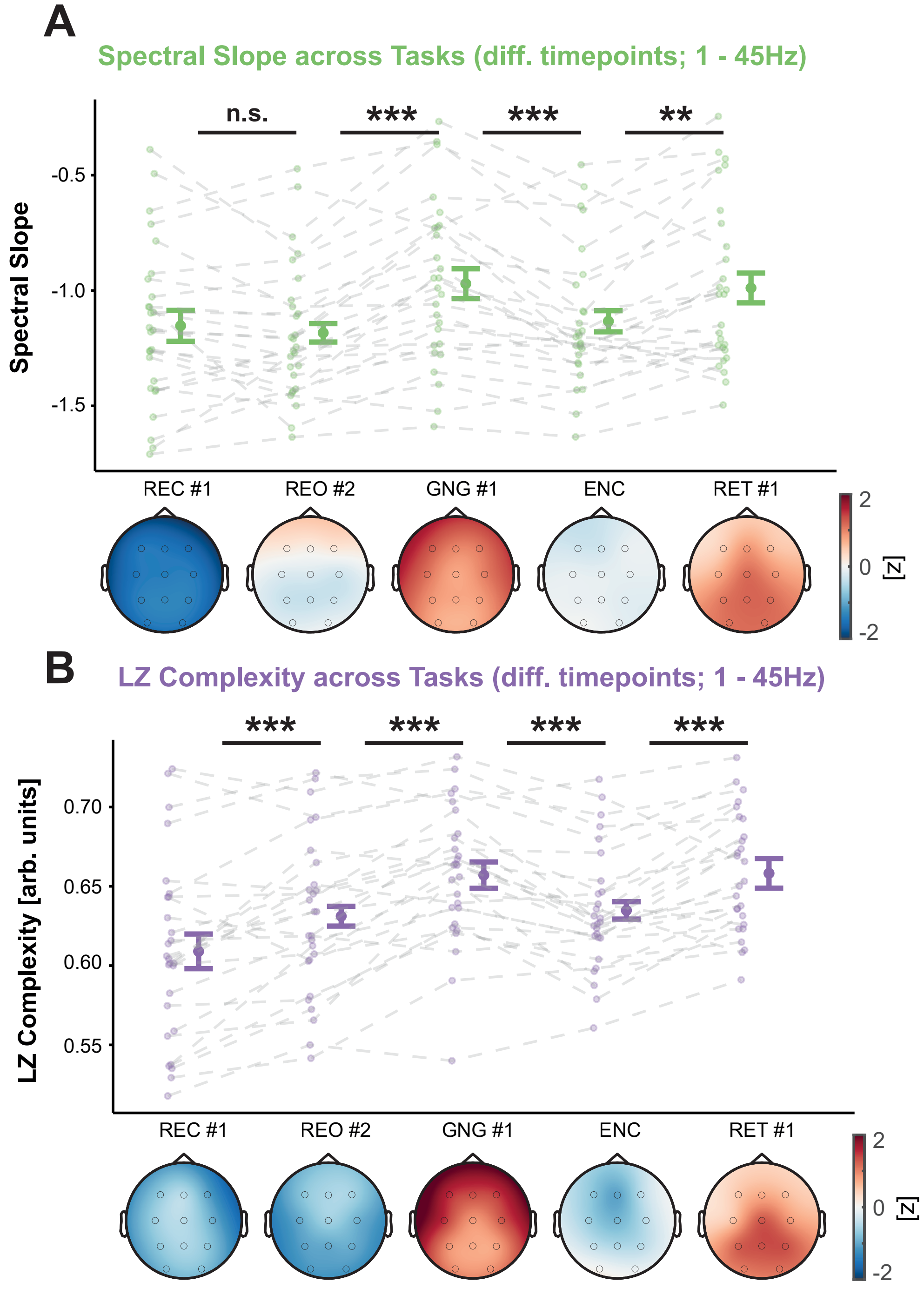

### Figure6_sup3.pdf

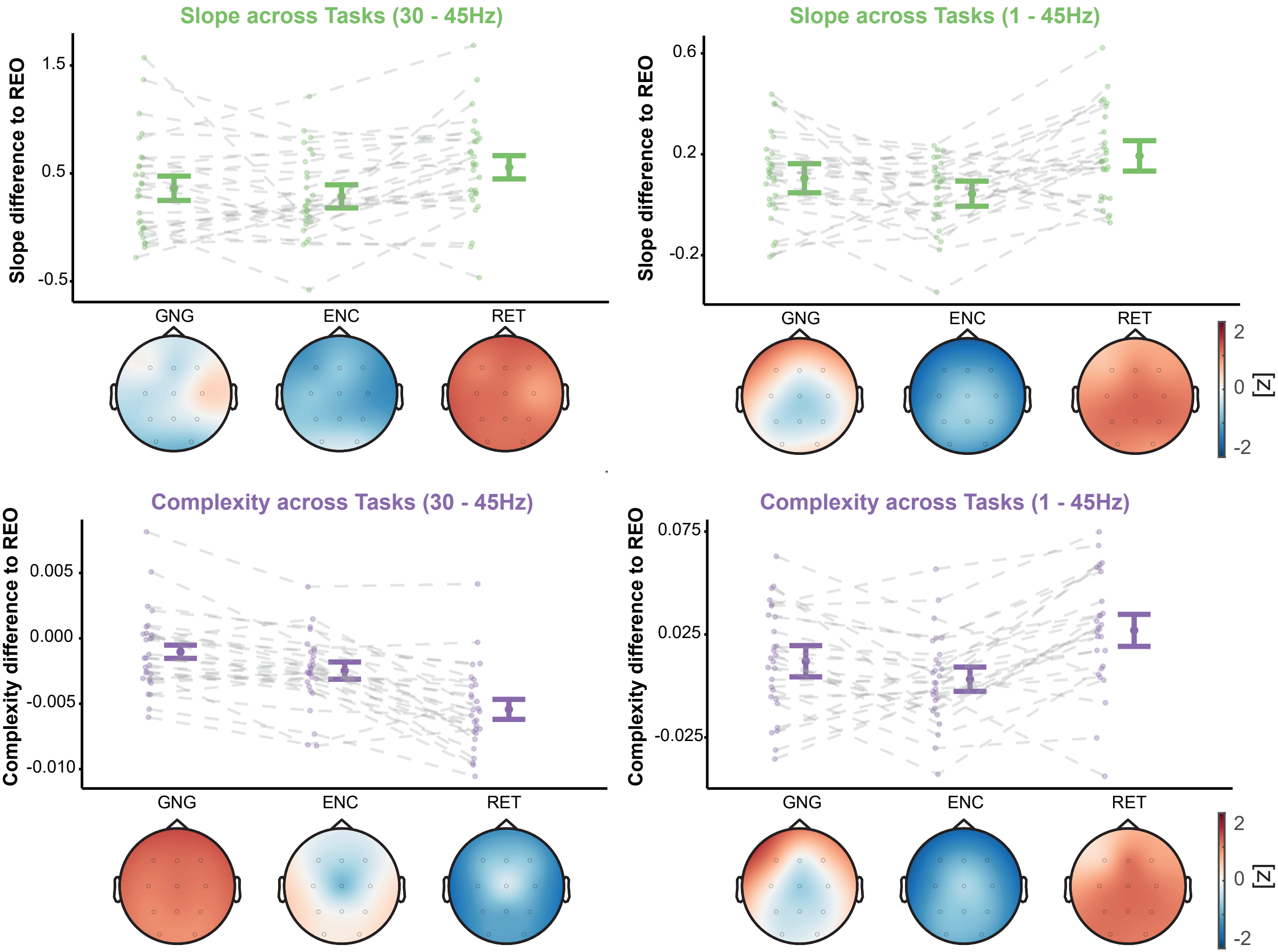

### Figure7_sup1.pdf

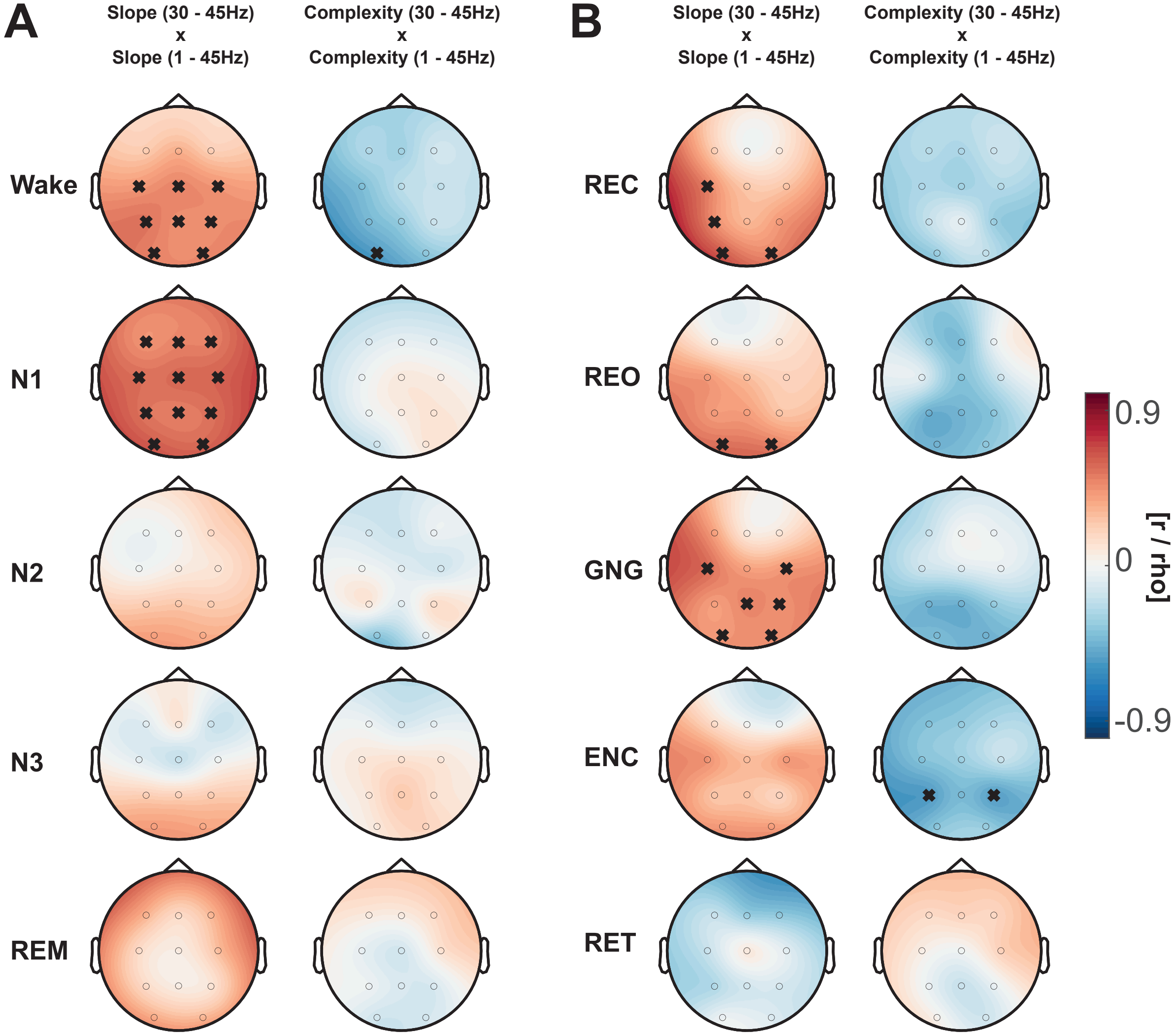

### Figure8_sup1.pdf

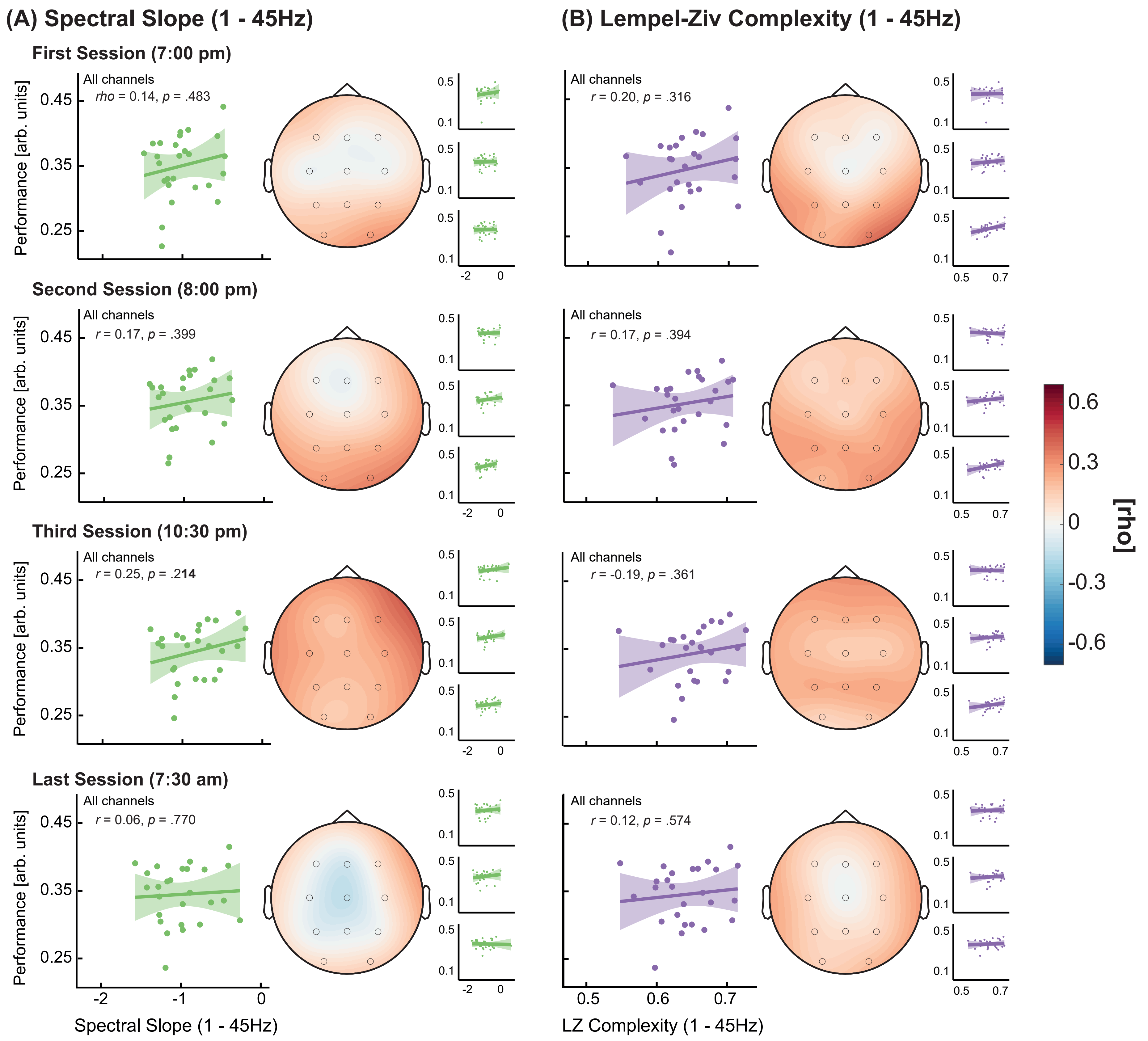

### Figure9_sup1.pdf

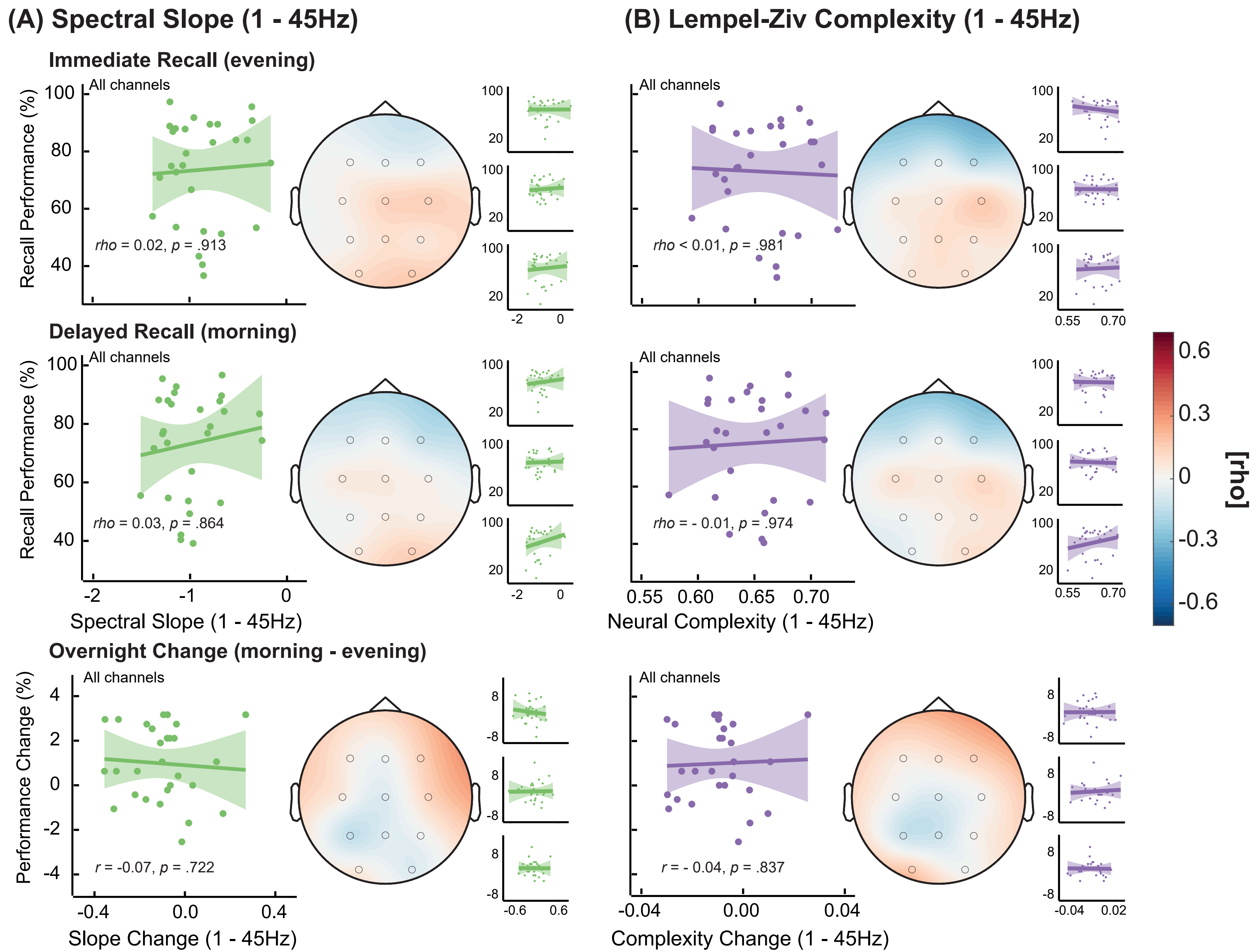
